# BDNF and TRiC-inspired Reagents Rescue Cortical Synaptic Deficits in a Mouse Model of Huntington’s Disease

**DOI:** 10.1101/2022.11.25.517123

**Authors:** Yingli Gu, Alexander Pope, Charlene Smith-Geater, Christopher Carmona, Aaron Johnstone, Linda Shi, Xuqiao Chen, Sarai Santos, Claire Cecile Bacon-Brenes, Thomas Shoff, Korbin M. Kleczko, Judith Frydman, Leslie M. Thompson, William C. Mobley, Chengbiao Wu

## Abstract

Huntington’s disease (HD) results from a CAG repeat expansion in the gene for Huntington (HTT) resulting in expansion of the polyglutamine (Q) tract in the mutant protein (mHTT). Synaptic changes are early manifestations of neuronal dysfunction in HD. However, the mechanism(s) by which mHTT impacts synapse formation and function is not well defined. Herein we explored HD pathogenesis in the BACHD and the ΔN17-BACHD mouse models of HD by examining cortical synapse formation and function in primary cultures maintained for up to 35 days (DIV35). We identified synapses by immunostaining with antibodies against pre-synaptic (Synapsin 1) and a post-synaptic (PSD95) marker. Consistent with earlier studies, cortical neurons from both WT and the HD models began to form synapses at DIV14; at this age there were no genotypic differences in synapse numbers. However, from DIV21 through DIV35 BACHD neurons showed progressively smaller numbers of synapses relative to WT neurons. Remarkably, BACHD synaptic deficits were completely rescued by treating cultures with BDNF. Building on earlier studies using reagents inspired by the chaperonin TRiC, we found that addition of the recombinant apical domain of CCT1 partially rescued synapse number. Unexpectedly, unlike BACHD cultures, synapses in ΔN17-BACHD cultures showed a progressive increase in number as compared to WT neurons, thus distinguishing synaptic changes in these HD models. Using multielectrode arrays, we discovered age-related functional deficits in BACHD cortical cultures with significant differences present by DIV28. As for synapse number, BDNF treatment prevented most synaptic deficits, including mean firing rate, spikes per burst, inter-burst interval, and synchrony. The apical domain of CCT1 showed similar, albeit less potent effects. These data are evidence that deficits in HD synapse number and function can be replicated *in vitro* and that treatment with either BDNF or a TRiC-inspired reagent can prevent them. Our findings support the use of cellular models to further explicate HD pathogenesis and its treatments.

## Introduction

Huntington’s disease (HD) a fatal disorder impacting movement and cognition, results from an expanded CAG repeat (>36) within the huntingtin gene, encoding a mutant Huntington protein (mHTT) with an expanded polyglutamine repeat (polyQ) (Group, 1993; Mangiarini et al., 1996; Sharp et al., 1995). HD is marked by severe cortical-striatal degeneration with clinical manifestations including chorea and other changes in movement, cognitive decline and neuropsychiatric disturbances(Finkbeiner, 2011; Lane et al., 2018; McColgan and Tabrizi, 2018{Finkbeiner, 2011 #3). The onset of HD symptoms is typically at 30 to 50 years of age; adolescent cases account for less than 10% (Frank, 2014). HD patients survive an average of 10-20 years following onset (Walker, 2007). Although pharmacological interventions have been approved for alleviation of symptoms(Caron et al., 2018; Novak and Tabrizi, 2011; Roos, 2010), treatments safe and effective for delaying the onset or reversing disease progression are yet to be discovered(Caron et al., 2018; Finkbeiner, 2011; Frank, 2014; Komatsu, 2021; Lane et al., 2018; Pan and Feigin, 2021).

Accumulation of mHTT is believed to mark disrupted protein homeostasis (Caron et al., 2018; Cepeda and Levine, 2022; Frank, 2014; Lane et al., 2018). How failed proteostasis may be causal for the many cellular deficits characteristic of HD is a topic of intense research interest. Indeed, mHTT is known to impact gene expression, mitochondrial function, endoplasmic reticulum (ER) function, protein turnover, axonal transport, and synaptic transmission (Bennett et al., 2007; Browne, 2008; Cepeda and Levine, 2022; Guo et al., 2016; Leitman et al., 2013; Li et al., 2003; Maity et al., 2022; Santos et al., 2010; Xie et al., 2010; Yoshii and Constantine-Paton, 2010). Defining the mechanism(s) by which mHTT contributes to pathogenesis promises to contribute significantly to understanding and treating HD.

A prominent clinical manifestation of HD is the structural and functional degeneration of cortical-striatal pathway. The degeneration of striatal medium spiny neurons (MSNs) is conjoined with loss of cortical pyramidal neurons (Vonsattel and DiFiglia, 1998). Increasing evidence links cortical-striatal atrophy and degeneration to deficiency in the levels or actions of brain-derived neurotrophic factor (BDNF)(Baydyuk and Xu, 2014; Gauthier et al., 2004; Hong et al., 2016; Su et al., 2014; Yu et al., 2018). BDNF is synthesized in many brain regions, albeit not in striatum (Conner et al., 1997; Hofer et al., 1990; Lindsay et al., 1994; Maisonpierre et al., 1990; Yan et al., 1997). BDNF, released from the cortex, binds to activate surface the receptor tyrosine kinase TrkB to provide critical trophic support to the striatum (Baydyuk and Xu, 2014; Ferrer et al., 2000; Lu et al., 2014; Yoshii and Constantine-Paton, 2010; Yu et al., 2018; Zhao et al., 2016; Zuccato and Cattaneo, 2007, 2009). Reduced BDNF/TrkB signaling is viewed as playing an important role in dysfunction and degeneration of MSNs and thus the cortical-striatal pathway in HD (Blumenstock and Dudanova, 2020; Cummings et al., 2007; Dallerac et al., 2011; Kovalenko et al., 2018; Milnerwood et al., 2006; Milnerwood and Raymond, 2007; Murphy et al., 2000; Nithianantharajah and Hannan, 2013; Puigdellivol et al., 2015). However, the underlying cellular mechanisms responsible for changes in BDNF delivery, release and signaling in striatum are yet fully understood. Deciphering the pathogenesis of HD will benefit from studies to explore the roles that BDNF plays in the formation and maintenance of cortico-striatal synapses.

Using microfluidic chambers to fluidically isolate cortical and striatal neurons to recreate the cortico-striatal circuit (Zhao et al., 2016), we have previously demonstrated that BACHD showed deficits in anterograde transport of BDNF and atrophy of striatal neurons. Importantly, we showed that BDNF release from cortical axons positively regulated the size of striatal neurons (Zhao et al., 2016). BACHD striatal neurons were atrophic when they were co-cultured with BACHD cortical neurons, but not with WT cortical neurons, pointing to reduced BDNF transport and release from BACHD cortical axons as causing atrophy. Our study thus argued for a presynaptic deficit in BDNF transport and release in BACHD neurons as responsible for striatal atrophy(Zhao et al., 2016).

To ask if failed proteostasis contributed to reduced BDNF transport and release we explored the impact of reagents derived from the chaperonin TRiC (Zhao et al., 2016). TRiC is a cytosolic chaperonin, comprised eight different subunits (CCT1-8) (Frydman, 2001; Frydman et al., 1992; Kalisman et al., 2012; Leitner et al., 2012) that contributes significantly to protein homeostasis and quality control (Gestaut et al., 2019; Roh et al., 2015; Thulasiraman et al., 1999). mHTT is a substrate of CCT (Kabir et al., 2011; Nollen et al., 2004) and increased expression of CCT prevents mHTT aggregation and toxicity (Kitamura et al., 2006; Rockabrand et al., 2007; Sontag et al., 2013; Tam et al., 2006; Tam et al., 2009). Conversely, reduction of CCT levels enhances mHTT aggregation(Kitamura et al., 2006; Rockabrand et al., 2007; Tam et al., 2006; Tam et al., 2009). Several studies have established that mHTT binds to the apical domain of subunit CCT1 (ApiCCT1) (Shen et al., 2016; Tam et al., 2006). Exogenous application of ApiCCT1 reduces aggregation of mHTT (Sontag et al., 2013) and remodels mHTT aggregates(Sahl et al., 2016). We discovered that ApiCCT1 and other TRiC-inspired reagents restored BDNF axonal trafficking in cortical neurons and through BDNF release prevented atrophy of striatal neurons in BACHD cortico-striatal circuit cultures. This evidence pointed to failed proteostasis as one mechanism by which BDNF deficits could contribute to HD pathogenesis(Zhao et al., 2016).

To further explore synapse deficits in HD, including a role for reduced BDNF release, herein we examined synapse formation and function in long-term cultures of E18 cortical neurons of WT mice and two mouse models: the BACHD(Gray et al., 2008) and the ΔN17-BACHD mouse in which the N-terminal first 17 residues of mHTT are deleted(Gu et al., 2015). Relative to BACHD mice, ΔN17-BACHD mice show more mHTT aggregation and earlier onset of neural deficits (Gray et al., 2008; Gu et al., 2015) and we hypothesized that ΔN17-BACHD neurons would demonstrate more severe synaptic deficits. Our results showed that both BACHD and ΔN17-BACHD cortical neurons formed the same number of synapses as WT neurons at DIV14 but that synapse numbers in BACHD cultures progressively decreased from DIV21 to DIV35. Remarkably, and unexpectedly, synapse numbers in ΔN17-BACHD cultures steadily increased and exceeded the number in WT cultures. BACHD cortical cultures also demonstrated age-related changes in synapse function with significant differences evident by DIV28. BDNF treatment prevented deficits in synapse number and function in BACHD cultures. ApiCCT1, while also effective, prevented only a subset of deficits. Taken together, the data are evidence for the ability to explore in vitro the synaptic pathogenesis of HD and for potent effects for BDNF in preventing changes in synapse number and function. They underscore the importance of studies to further explore the role played by BDNF in HD pathogenesis and treatment.

## Materials and Methods

### Ethical Statement

All experiments involving the use of laboratory animals have been approved by the Institutional Animal Care and Use Committee of University of California San Diego. Surgical and animal procedures were carried out strictly following the NIH Guide for the Care and Use of Laboratory Animals.

### Reagents and antibodies

10x Hanks’ Balanced Salt solution (HBSS), 2.5% Trypsin-EDTA (10x), 100x Penicillin-Streptomycin, 100x Glutamax, 50x B27, Neuronal basal media were all purchased from Invitrogen. DNase I grade II from bovine pancreas (Roche, Cat#10104159001) was dissolved in 1x HBSS at 10 mg/ml (10x) and filter-sterilized. FBS was from Mediatech Inc (Cat# 35-010-CV). 0.1% Poly-L-Lysine (Cultrex® Poly-L-Lysine) was from Trevigen (Gaithersburg, MD; Cat# 3438-100-01). Human recombinant BDNF was from Genentech (San Francisco, CA). Hoechst 33258 was from Sigma (Cat#861405). All other chemicals and lab wares were from Bio-Rad, Fisher, Sigma, VWR.

Mouse anti-PSD95 (Clone 28/47 from Biolegend, Cat. #810401), rabbit monoclonal anti-Synapsin I (Cell Signaling, Cat. #D12G5), mouse anti-synaptobrevin 2 (104211) (SYSY; 1:5000 dilution), mouse anti-synaptophysin (PA1-1043; 1:5000 dilution), mouse anti-synaptotagmin (610434; 1:2000 dilution) (BD), mouse anti-SNAP25 (ADI-VAM-SV012; 1:2000 dilution) (Enzo), mouse anti-β-actin was from Santa Cruz Biotechnology (Cat. #47778, 1:2000 dilution). Secondary goat anti-rabbit, goat anti-mouse antibodies conjugated to Alexa-488, 568 were from Invitrogen. All antibodies were used at dilutions following manufacture’s instructions. Paraformaldehyde was purchased from Sigma.

### Neuronal culture and maintenance

Established protocols were followed to set up cortical neurons collected from mouse E18 embryos(Fang et al., 2017; Zhao et al., 2016). Briefly, cortical tissues were extracted from E18 mouse embryos and extensively rinsed in HBSS with 1% Penicillin-Streptomycin, followed by dissociation in 0.25% trypsin with 1 mg/ml DNase I. Cortical neurons were isolated and plated with plating media (Neurobasal with 10% FBS, 1xB27,1xGlutaMAX) onto either glass coverslips for immunostaining or into 12 well plates for biochemistry at appropriate density. Both the coverglasses and plates were pre-coated with poly-L-lysine (Invitrogen). Plating medium was replaced with maintenance medium (Neurobasal, 1xB27, 1xGlutamax) the following day. Only 2/3 of the media was replaced every other day until conclusion of the experiments.

### Immunostaining and confocal microscopy

E18 cortical neurons from wild type (WT), BACHD and ΔN17 BACHD mice were cultured on coverglasses. The coverglasses were precoated with 0.1% poly-D-Lysine (1 hr at RT) followed by three rinses in sterile dH_2_O and air dry. Cultures at DIV7 (Days-In-Vitro), DIV14, DIV21, DIV28 and DIV35 were rinsed and were fixed in 4% paraformaldehyde for 15 mins at RT. The samples were washed 3 times with PBS and were blocked and permeabilized in 5% goat serum containing 0.2% TritonX-100 for 15 mins at RT. Neurons were first incubated with a mouse anti-PSD95 antibody (1:200 dilution in PBS) for 3 hrs at RT. The samples were washed 3 times (5 min each) in PBS and were incubated with the rabbit monoclonal antibody against Synapsin I (1:200 dilution in PBS) at 4℃ overnight. Samples were washed in PBS 3 times (5 min each) and were incubated with goat anti-mouse Alexa 488 and goat anti-rabbit Alexa 568 secondary antibody conjugates (both at 1:600 dilution in PBS) for 1hr at RT. Hoechst 33258 (1.0 µg/ml in PBS) was used to stain nuclei for 5 min at RT. Samples were washed and mounted onto slides. Images of neurons were captured with a Leica SP confocal microscope under a 63x oil objective lens with a 1.6x zoom factor.

We followed published methods (Paul et al., 2013; Rizk et al., 2014) to quantify synaptic formation of cortical neurons of WT and BACHD. Segmentation was done on the neurons to identify puncta of PSD95 and Synapsin I by using squassh in the NIH MOSAIC image suite(Paul et al., 2013; Rizk et al., 2014). Pearson’s colocalization co-efficient (PCC) between PSD95 and Synapsin I, percentile of PSD95 signals overlapping with Synapsin I signals, percentile of Synapsin I signals overlapping with PSD95 signals were analyzed and quantitated using the NIH ImageJ MOSAIC Suite. ImageJ was also used to measure the soma size and the size of PSD95, Synapsin I puncta. We also quantitated synaptic density analysis normalized against the length of neurites per 100μm.

### BDNF treatment

To test a rescuing effect of synaptic deficits by BDNF, exogenous BDNF was included in the neuronal maintenance media. WT or BACHD cortical neuronal cultures were treated with media containing either BDNF (100 ng/ml, final concentration) or vehicle starting at either DIV7 or DIV14 up till to DIV21. Neurons were thus treated for either 14 days or 7 days respectively. Synaptic staining (PSD95/Synapsin I) and quantitation of PCCs were carried out at DIV21 as described above.

### ApiCCT1 Treatment

Recombinant ApiCCT1 was purified as previously described (Shen et al., 2016). Prior to addition to the culture media, the ApiCCT1 protein preparations were desalted and reconstituted into Neurobasal media with the use of Zeba™ Spin Desalting Columns, 7K MWCO, 0.5 mL (ThermoFisher, Cat# 89882). Re-purified ApiCCT1 was added to the culture media at a final concentration of 0.1μM at DIV14. The media were changed with fresh ApiCCT1 every other day prior to analysis at DIV21.

### BDNF ELISA

Rapid ELISA Kits for measuring mature BDNF (Cat# BEK-2211-2P) were purchased from Biosensis Pty Ltd (51 West Thebarton Road, Thebarton, South Australia). To measure BDNF secretion, conditioned media were collected from WT and BACHD cortical neuronal cultures. The media (∼400 μl) were lyophilized overnight, reconstituted with 120 μl sterile dH_2_O by vigorous vortex at RT. Samples were centrifuged at 14k rpm at 4°C for 10 min. 100 μl clear supernatants were removed and the amounts of BDNF were measured and quantified against the standard curve following the manufacture’s instruction. To normalize BDNF secretion on a total protein base, we collected neurons from each well, solubilized in radio-immunoprecipitation assay (RIPA) buffer and centrifuged to produce the supernatants. The protein concentrations were determined by BCA method using a NanoDrop™ 2000/2000c Spectrophotometer (ThermoFisher) and the results were used to normalize the levels of BDNF in the secreted media.

### Multielectrode array (MEA)

Neurons were plated on Poly-D-Lysine coated (100 µg/ml – Sigma Aldrich) CytoView MEA 24 well plates (Axion Biosystems) at a density of 100,000 cells per well, plated in 10 µl of plating medium. These were allowed to attach at 37 ℃, 5% CO_2_ for 30-45 minutes until the neurons settled, at which time the wells were flooded with 500 µl of plating medium in 2 increments of 250 µl each. The medium was changed to maintenance medium after 24 hours and fed every 48 hours, performing a 2/3 medium change each time. At DIV14, the first recording on the Maestro Edge was performed for 10 minutes; recordings were acquired using the AxIS Navigator Software v3.2.3.1. After recording, the medium was changed and included BDNF (50ng/ml), ApiCCT1 (0.1µM) or Vehicle control prepped as previously described (PMID: 30955883). A final recording was performed as above at DIV 28. Analysis was performed by running Batch process in AxIS Navigator v2.0.4.21, “re-recording” data for consistency at the end of each experiment. For spike detection, the adaptive threshold crossing was used. The burst detection setting had a maximum inter spike interval of 100 ms and minimum of 5 spikes/burst required. For network statistics, the minimum number of spikes required was 30 with at least 35% of the well electrodes participating in the network activity.

### Western blot analysis and quantification

Primary cortical neurons from BACHD and WT littermates were cultured and collected at DIV21, DIV28. After lysis in RIPA buffer containing PMSF protease inhibitor on ice for 30 min, the cell lysate was centrifuged at 13,300 rpm at 4℃ for 10 min. The supernatants were collected and SDS loading buffer added. The protein samples were denatured at 95℃ for 4 min and loaded onto 10% of SDS-PAGE (sodium dodecyl sulfate polyacrylamide gels) for separation. Separated proteins were electro-transferred onto cellulose nitrate membranes, blocked in 5% skim milk diluted in TBST at RT for 1 hr. And the membranes were incubated with: mouse primary antibodies against synaptophysin (Synaptic Systems GmbH, 1:1,000), synaptobrevin (mouse, 1:5,000), synaptotagmin (mouse, 1:2,000), SNAP25 (mouse, 1:5,000), PSD95 (mouse, 1:1,000) and β-actin (C4) (mouse, sc-47778 from Sant Cruz) at 4℃ overnight, and then incubated in corresponding secondary antibodies (goat anti-mouse, 1:5,000 or 1:10,000). The blots were washed and developed in ECL-Clarity (BioRad). The blots were imaged using ChemiDoc XRS+ (Bio-Rad) and quantitated using ImageLab 6.0.1 software (BioRad). Protein levels were expressed as the ratio of each immunoreactive band and the levels of β-actin.

### Statistical analysis

Significance analysis was carried out using Prism. Significances were calculated using either unpaired t-test (Mann-Whitney), One Way ANOVA (Dunnett’s post-test) or Two Way ANONA (Tukey’s multiple comparison). These methods are specifically indicated in relevant Figures and Tables. Values are the mean ± SEM of three different and independent experiments. Ns: non significance; * p<0.05; **p<0.01; *** p<0.001; **** p<0.0001.

## Results

### Age-related deficits in cortical neuron synapse number in the BACHD, but not in the ΔN17 BACHD model

We used two HD mouse models to explore synapse formation and function (Gray et al., 2008; Gu et al., 2015). The BACHD transgenic model expresses the full-length human mutant huntingtin (mHTT) gene (polyQ97) using a bacterial artificial chromosome (BAC); the mutant transgene is expressed at a level ∼1.5-2.0 fold the wildtype allele in BACHD mice(Gray et al., 2008). The ΔN17-BACHD model is based on BACHD except that the N-terminal 2-16-amino-acid (N17) residues were deleted (Gu et al., 2015); the transgene is expressed at a similar level to the transgene allele in BACHD mice. Compared to the BADHD model, ΔN17-BACHD mice develop more mHtt aggregates and show more severe neuronal deficits at early stages (Gray et al., 2008; Gu et al., 2015). Based on observations from our lab and others(Gray et al., 2008; Zhao et al., 2016), we predicted deficits in synapse formation and function in cortical neuron cultures in the BACHD mouse and that they would be more severe in ΔN17-BACHD model. To test for differences in synapse formation we cultured E18 cortical neurons from BACHD, ΔN17-BACHD and WT littermate controls. Cultures were maintained for up to 35 days in vitro (DIV35). Cultures at DIV14, 21, 28, 35 were analyzed for synapse number to report on the combined contributions of synapse formation and maintenance. Synapses were quantified by immunostaining with antibodies against Synapsin I for pre-synapses and against postsynaptic density 95 (PSD95) for post-synapses. Images were collected using confocal microscopy. The raw images were evaluated using ImageJ /MOSAIC Suite. For each channel, the background values were subtracted and the images were then segmented using the Squassh function to isolate puncta. We captured several parameters relevant to synapses at each DIV: 1) colocalization between these markers, denoted and quantitated as Pearson’s Colocalization Co-efficient (PCC); and 2) the percentage of post- and pre-synaptic staining that contributed to the count of synapses, as revealed by assessing the extent to which PSD95 signals overlapped with Synapsin I (i.e. % of PSD95/Synapsin I) and conversely, Synapsin I signals that overlapped with PSD95 (% of Synapsin I/PSD95). Representative images and corresponding quantitative results are presented in Fig. 1 through 4 for DIV14 through DIV35. In addition, we evaluated the size of Synapsin1 and PSD95 puncta and synapse density (Fig. 5).

**Figure 1.**
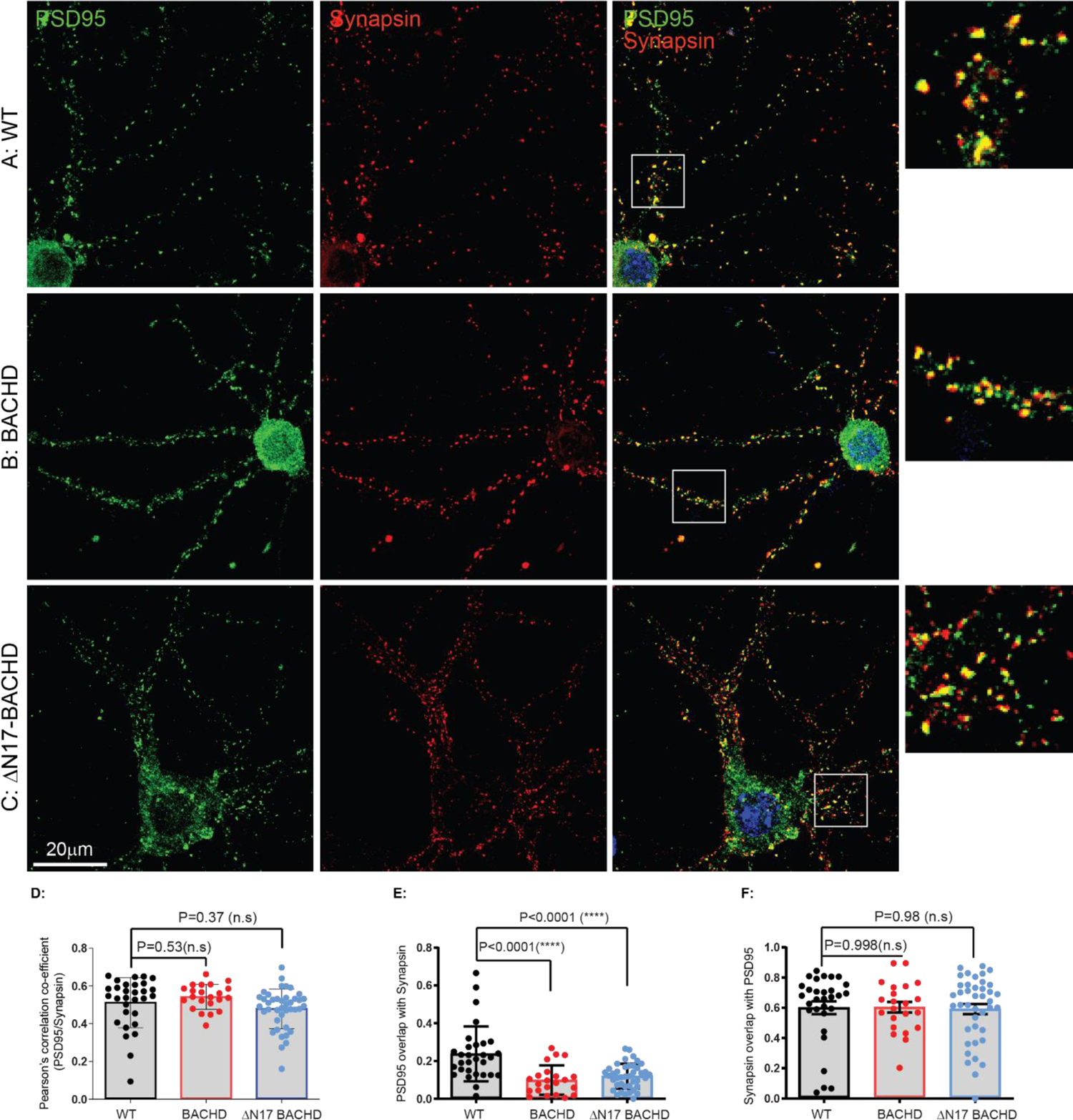
Synaptic analysis of cortical neurons from WT, BACHD and ΔN17-BACHD at DIV14. E18 cortical neurons from WT, BACHD, ΔN17-BACHD were dissected, cultured on PLL-coated cover-glasses and maintained as described in Materials and Methods. At DIV14, a set of samples from each genotype were fixed and immunostained with a rabbit monoclonal antibody against synapsin I (1/200) and a mouse monoclonal against PSD95 (1/200). Nuclei were stained with Hoechst (1:10000). The images were captured under a 63x oil objective using a Leica SP6 confocal microscope. Representative images are shown and co-localization between synapsin I and PSD95 was quantitated using ImageJ Suite Plugin. (A) (B) (C) show representative images of PSD95 (green) and Synapsin I (red) staining respectively in WT, BACHD, ΔN17-BACHD cortical neurons. Regions of interest marked by white boxes are magnified and shown on the right. (D) Comparison of post- and presynaptic marker colocalization using Pearson’s Colocalization Coefficients (PCC). (E) Analysis of PSD95 signals that overlapped with Synapsin I. (F) Analysis of Synapsin I signals that overlapped with PSD95. Results are shown as mean ± SEM. The numbers of images were analyzed are: n=30 for WT, n=22 for BACHD, n=41 for ΔN17-BACHD. The data represent at least 4-5 independent cultures. Significance analysis was carried out using Prism. Statistical significances were calculated by Sidak’s multiple comparisons test of One-Way ANOVA. n.s.= non significance. All p values are shown in the graphs.

### Synapse numbers differed by genotypes after DIV14

Synapses were readily evident by DIV14 in cultures from WT, BACHD, and ΔN17-BACHD mice, a finding consistent with reports demonstrating that neurons form synapses beginning at DIV6-7, with fully functional synapses appearing at DIV12-14(Kavalali et al., 1999; Nosyreva et al., 2013). Analysis of synapse number, as reflected in PCC, at each time point showed the following for WT neurons: cortical neuron synapse number was stable from DIV14 (0.511 ± 0.024) (Fig. 1A, D) to DIV21(0.565 ± 0.010, p>0.05 as compared to DIV14) (Fig. 2A, D) and to DIV28 (0.549 ± 0.016, p>0,05 as compared to DIV14) (Fig. 3A, D). Between DIV28 to DIV35 the PCC value was slightly increased to 0.620 ± 0.026, p<0,05 as compared to DIV14 (Fig. 4A, D); statistically (One-Way ANOVA with Dunnett’s Multiple Comparison Test), there was no difference across ages in WT culture (p>0.05 DIV21 vs DIV14; p>0.05 DIV28 vs DIV14) except at DIV35 (p<0.01, DIV35 vs DIV14).

**Figure 2.** Synaptic analysis of cortical neurons from WT, BACHD and ΔN17-BACHD at DIV21. Neuronal culture, immunostaining and quantitation are as described in Fig 1. (A) (B) (C) show representative images of PSD95 (green) and Synapsin I (red) staining respectively in WT, BACHD, ΔN17-BACHD cortical neurons. Regions of interest marked by white boxes are magnified and shown on the right. (D) Comparison of post- and presynaptic marker colocalization using Pearson’s Colocalization Coefficients (PCC). (E) Analysis of PSD95 signals that overlapped with Synapsin I. (F) Analysis of Synapsin I signals that overlapped with PSD95. Results are shown as mean ± SEM. The numbers of images were analyzed are: n=65 for WT, n=45 for BACHD, n=58 for ΔN17-BACHD. The data represent at least 4-5 independent cultures. Significance analysis was carried out using Prism. Statistical significances were calculated by Sidak’s multiple comparisons test of One-Way ANOVA. n.s.= non significance. All p-values are shown in the graphs.

**Figure 3.**
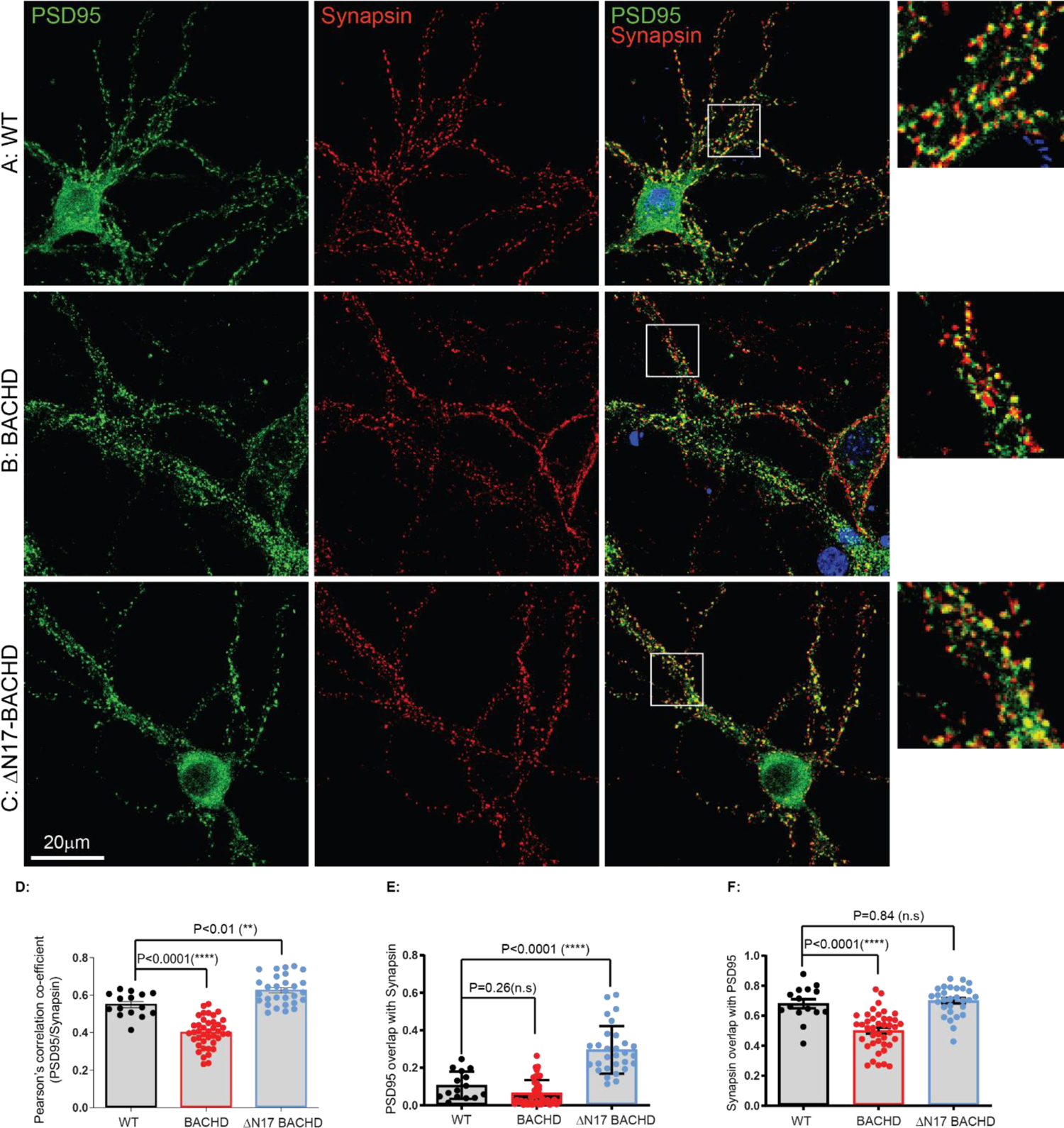
Synaptic analysis of cortical neurons from WT, BACHD and ΔN17-BACHD at DIV28.0 Neuronal culture, immunostaining and quantitation are described as in Fig 1. Representative images are shown and co-localization between Synapsin I and PSD95 was quantitated using ImageJ Suite Plugin. (A) (B) (C) show representative images of PSD95 (green) and Synapsin I (red) staining respectively in WT, BACHD, ΔN17-BACHD cortical neurons. White boxes demarcating regions of interest magnified on the right. (D) Comparison of post- and presynaptic marker colocalization using Pearson’s Colocalization Coefficients (PCC). (E) Analysis of PSD95 signals that overlapped with Synapsin I. (F) Analysis of Synapsin I signals that overlapped with PSD95. Results are shown as mean ± SEM. The numbers of images were analyzed for n=15 (WT), n=41 for BACHD, for ΔN17-BACHD n=29. The data represent at least 4-5 independent cultures. Significance analysis was carried out using Prism. Statistical significances were calculated by Sidak’s multiple comparisons test of One-Way ANOVA. n.s.= non significance. All p values are shown in the graphs.

**Figure 4.**
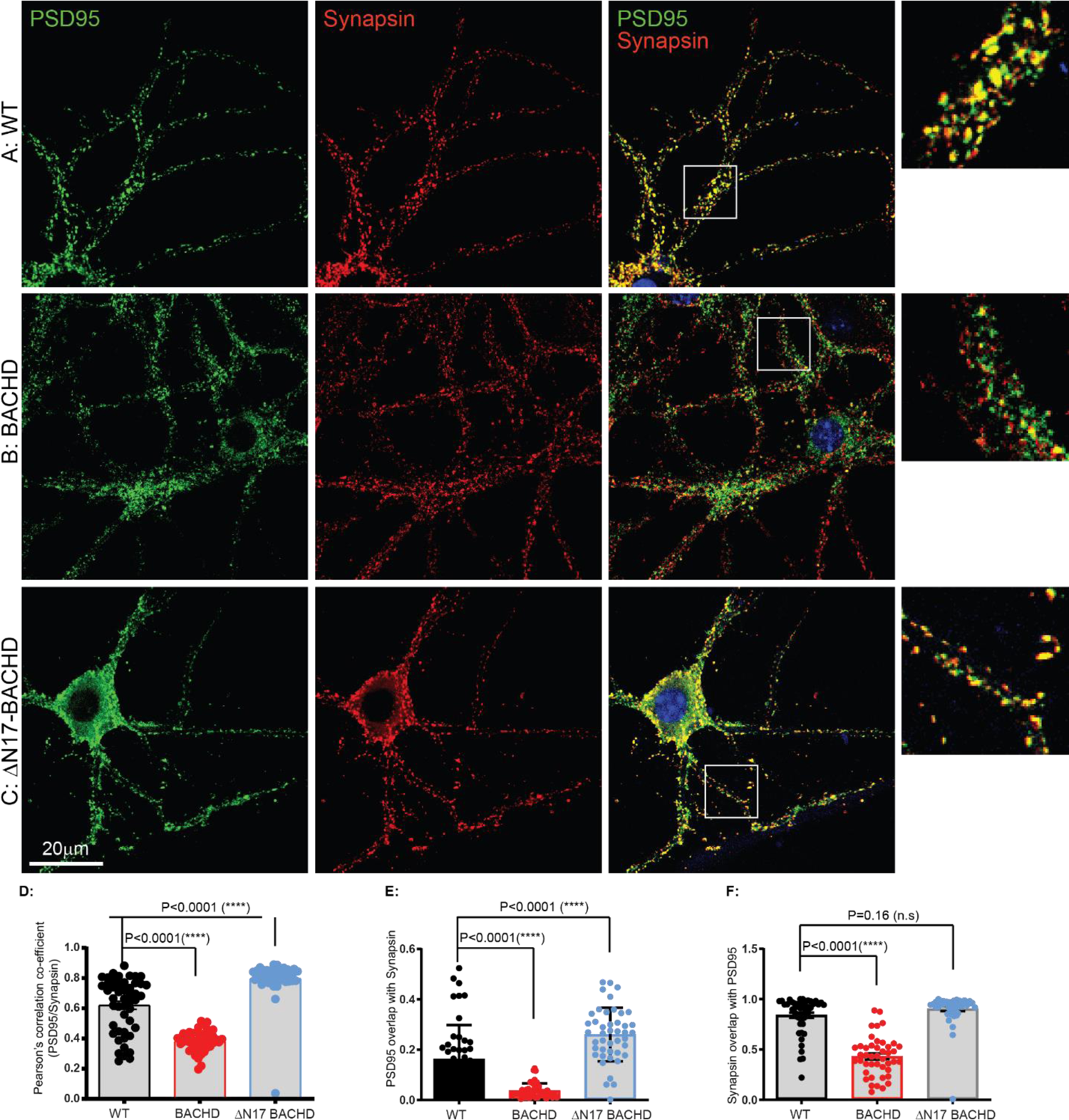
Synaptic analysis of cortical neurons from WT, BACHD and ΔN17-BACHD at DIV35. E18 cortical neurons from WT, BACHD, ΔN17-BACHD were dissected, cultured on PLL-coated cover-glasses and maintained as described in Materials and Methods. At DIV35, a set of samples from each genotype were fixed and immunostained with a rabbit monoclonal antibody against synapsin I (1/200) and a mouse monoclonal against PSD95 (1/200). Nuclei were stained with Hoechst (1:10000). The images were captured under a 63x oil objective using a Leica SP6 confocal microscope. Representative images are shown and co-localization between synapsin I and PSD95 was quantitated using ImageJ Suite Plugin. (A) (B) (C) show representative images of PSD95 (green) and Synapsin I (red) staining respectively in WT, BACHD, ΔN17-BACHD cortical neurons. Regions of interest marked by white boxes are magnified and shown on the right. (D) Comparison of post- and presynaptic marker colocalization using Pearson’s Colocalization Coefficients (PCC). (E) Analysis of PSD95 signals that overlapped with Synapsin I. (F) Analysis of Synapsin I signals that overlapped with PSD95. Results are shown as mean ± SEM. The numbers of images were analyzed are: n=48 (WT), n=44 (BACHD), n=44 (ΔN17-BACHD). The data represent at least 4-5 independent cultures. Significance analysis was carried out using Prism. Statistical significances were calculated by Sidak’s multiple comparisons test of One-Way ANOVA. n.s.= non significance. All p values are shown in the graphs.

Analysis of BACHD neurons showed: cortical neuron synapse number was equivalent to WT neurons at DIV14 (0.542 ± 0.014) (p=0.530) (Fig. 1B, D). However, by DIV21 (0.422 ± 0.019) (Fig. 2B, D) synapse number decreased, a change that progressed through DIV28 (0.400 ± 0.012) (Fig. 3B, D) (p<0.0001) and DIV35 (0.381 ± 0.010) (Fig. 4B, D) (p<0.0001); relative to BACHD neurons at DIV14, synapses at all subsequent DIVs were statistically significantly different (p<0.001). Analysis of ΔN17-BACHD neuron cultures showed: cortical neuron synapse number was not significantly different from WT at DIV14 (0.479 ± 0.016)(p=0.368) (Fig. 1C, D) or DIV21 (0.560 ± 0.017)(p=0.971) (Fig. 2C, D), but was significantly increased at DIV28 (0.626 ± 0.014)(p=0.004) (Fig. 3C, D) and increased further at DIV35 (0.796 ± 0.019)(p<0.0001) (Fig. 4C, D); relative to ΔN17-BACHD neurons at DIV14, synapses at all subsequent DIVs were statistically significant(p≤0.01).

The changes in PCC for WT, BACHD and ΔN17-BACHD neurons from DIV14-35 are plotted in Fig. 5A (Also See **Table 1**). We conclude that while BACHD and ΔN17-BACHD cortical neuron synapses were equivalent in number to those in WT cultures at DIV14, they differed thereafter with BACHD showing progressively fewer synapses and ΔN17-BACHD neurons displaying progressively more synapses.

**Figure 5.**
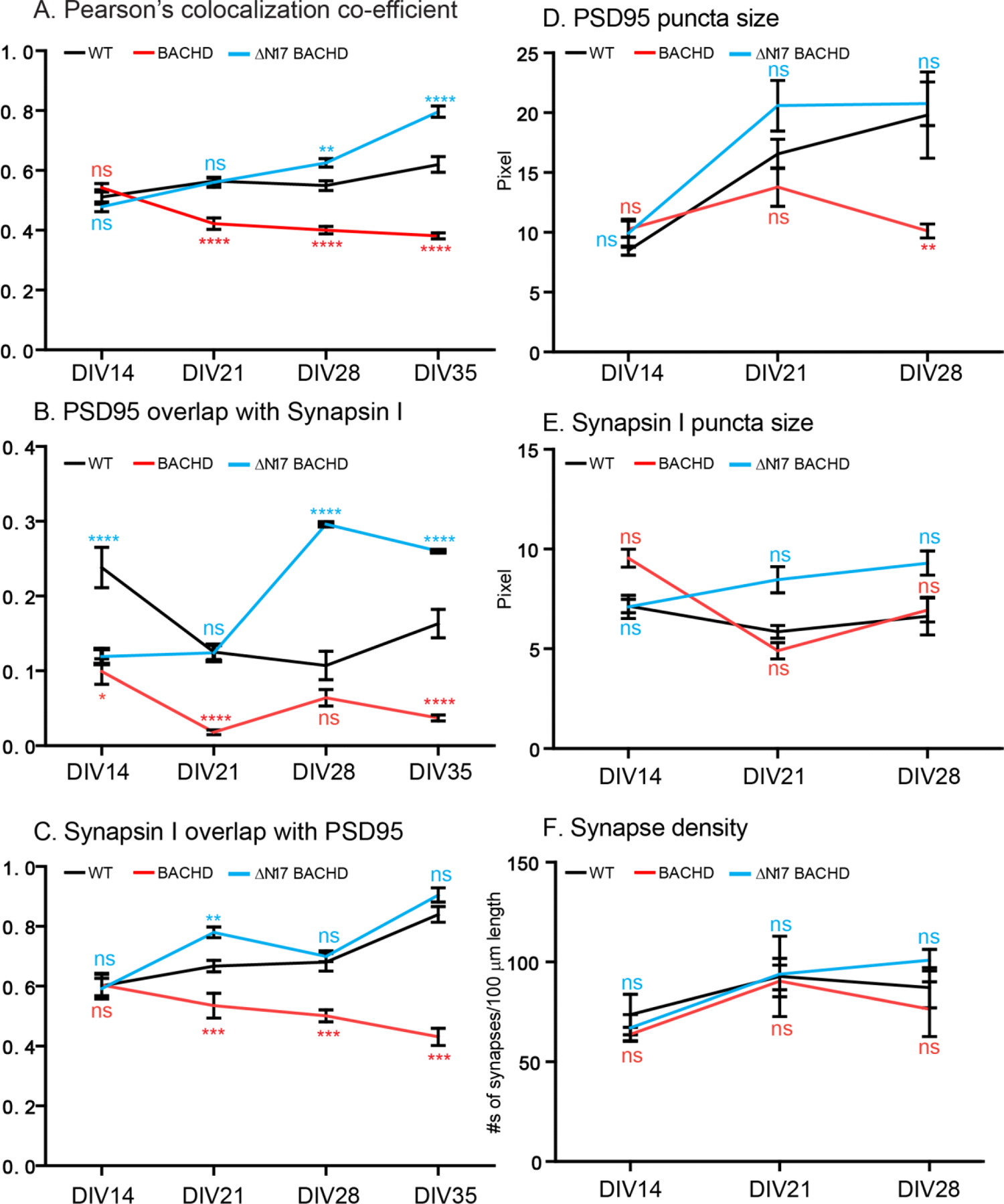
Time course of synaptic formation of cortical neurons from WT, BACHD and ΔN17-BACHD from DIV14-DIV35. A time course study of Pearson’s colocalization co-efficient values from DIV14 to DIV 35 (in Figure 1-4) was plotted to show the progression of synaptic formation in WT, BACHD and ΔN17-BACHD cortical neurons. (A) Time-dependent changes of Pearson’s Colocalization Coefficients. (B) Time-dependent changes of PSD95 signals that overlapped with Synapsin I. (C) Time-dependent changes of Synapsin I signals that overlapped with PSD95. (D) Measurements for PSD95 puncta size in WT, BACHD and ΔN17-BACHD cortical neurons from DIV14 to DIV28. (E) Measurements for Synapsin I puncta size in three genotypes from DIV14-DIV28. (F) Measurements for synaptic density in three genotypes from DIV14-28. Significance analysis was carried out using Prism. Statistical significances were calculated by One-Way ANOVA. n.s.= non significance. \**P* < 0.05, \*\**P* < 0.01, \*\*\**P* < 0.001, \*\*\*\**P* < 0.0001.

**Table 1.** Analysis of synaptic formation in cultured neurons of WT, BACHD, ΔN17 BACHD (I)

|  | Pearson's Colocalization Co-efficient (PCC) |  |  | % of PSD95/Synapsin I |  |  | % of Synapsin I/PSD95 |  |  |
| --- | --- | --- | --- | --- | --- | --- | --- | --- | --- |
| | WT | BACHD | ΔN17 BACHD | WT | BACHD | ΔN17 BACHD | WT | BACHD | ΔN17 BACHD |
| DIV14 | 0.511±0.024 | 0.542±0.014<sup>ns</sup> | 0.479±0.016<sup>ns</sup> | 0.238±0.027 | 0.099±0.017<sup>****</sup> | 0.119±0.010<sup>****</sup> | 0.601±0.042 | 0.604±0.036<sup>ns</sup> | 0.592±0.034<sup>ns</sup> |
| DIV21 | 0.565±0.010 | 0.422±0.019<sup>****</sup> | 0.560±0.017<sup>ns</sup> | 0.125±0.010 | 0.018±0.003<sup>****</sup> | 0.124±0.011<sup>ns</sup> | 0.667±0.019 | 0.535±0.041<sup>****</sup> | 0.78±0.018<sup>**</sup> |
| DIV28 | 0.549±0.016 | 0.4±0.012<sup>****</sup> | 0.626±0.014<sup>**</sup> | 0.107±0.019 | 0.064±0.011<sup>ns</sup> | 0.296±0.024<sup>****</sup> | 0.68±0.03 | 0.501±0.02<sup>****</sup> | 0.7±0.018<sup>ns</sup> |
| DIV35 | 0.620±0.026 | 0.381±0.01<sup>****</sup> | 0.796±0.019<sup>****</sup> | 0.163±0.019 | 0.037±0.004<sup>****</sup> | 0.26±0.016<sup>****</sup> | 0.84±0.026 | 0.431±0.029<sup>****</sup> | 0.905±0.024<sup>ns</sup> |

Significance test: One Way Anova (Dunnett's post test). ns= non-significance, p>0.05; \* p<0.05; \*\* p<0.01; \*\*\* p<0.001; \*\*\*\* p<0.0001

### Additional synaptic parameters distinguish WT from both BACHD and ΔN17-BACHD

To further analyze synapses, we examined additional synaptic parameters. First, we quantitated the percent to which was PSD95 colocalized with Synapsin 1 (i.e. PSD 95/Synapsin I) and the percent of Synapsin 1 colocalized with PSD95 (i.e. Synapsin 1/PSD95). While the number of synapses did not distinguish genotypes at DIV14, there was a difference in PSD95/Synapsin 1 at this age (Figure 1 E, **Table 1**), pointing to differences relative to WT even at this early age in the HD models. We found in WT cultures that PSD 95/Synapsin I was highest at DIV14 (0.238 ± 0.027) (Fig. 1A, E), with lower values at DIV21 (0.125 ± 0.010) (p<0.0001) (Fig. 2A, E), DIV28 (0.107 ± 0.019) (p=0.002) (Fig. 3A, E) and DIV35 (0.163±0.019) (p=0.029) (Fig. 4A, E). The overall reduction over time in the ratio of PSD95/Synapsin 1 was significant (p<0.0001). In the same cultures, WT neurons showed a gradual increase in Synapsin 1/PSD95 from DIV14 (0.601 ± 0.042) (Fig. 1A, F) to DIV21 (0.667 ± 0.019) (p=0.322) (Fig. 2A, F), DIV28 (0.680 ± 0.030) (p=0.480) (Fig. 3A, F), and DIV35 (0.840 ± 0.026) (p<0.0001) (Fig. 4A, F). The overall increase over time in the ratio of Synapsin 1/PSD95 was significant (p<0.0001). The changes in the overlap of PSD95 and Synapsin 1 signals could reflects changes at the pre-, post-synapses, or both. To differentiate between these possibilities, we examined with aging the size of PSD95 and Synapsin 1 puncta as well as synapse density. There was an increase in the size of PSD95 puncta, with relative stability in the size of Synapsin 1 puncta, together with an increase in synapse density (Fig. 5D, E, F). These data suggest that an increase in the size or number of postsynaptic complexes resulted in an increase in synapse density.

In BACHD cultures the percent for PSD95/Synapsin I was lower than in WT cultures at DIV 21 onward, significantly so at DIV 21 and 35 (**Table 1**). The values relative to WT were as follows: at DIV14 (Fig. 1E), (BACHD vs WT: 0.099±0.017 vs 0.238±0.027; p<0.0001); at DIV 21 (Fig. 2E) (0.018±0.003 vs 0.125± 0.01; p<0.0001); at DIV28 (Fig. 3E) (0.064±0.011 vs 0.107±0.019; p>0.05; and at DIV35 (Fig. 4E) (0.037±0.004 vs 0.163±0.019; p<0.0001). The overall reduction over time in the ratio of PSD95/Synapsin 1 in BACHD cultures was significant (p<0.0001). Synapsin 1 overlap with PSD95 was also decreased from DIV 21 onward. The values were_0.604±0.036for DIV14; 0.535±0.041 for DIV21; 0.501±0.02 for DIV28; 0.431±0.029 for DIV35). The overall reduction over time in the ratio of Synapsin 1/PSD95 was significant (p<0.0001). During the period from DIV14 to 28, the size of PSD95 puncta were smaller in BACHD than in WT cultures (Fig 5D, **Table 2**), at DIV 28 the size of PSD95 puncta was significantly smaller (WT:19.9±3.61; BACHD:10.11±0.59; p<0.0057) while the size of Synapsin 1 puncta was not different (WT: 6.64±0.94; BACHD:6.95±0.6; p=0.9826) (Fig. 5E and **Table 2**). These synaptic parameters were correlated with a decrease in synapse density between DIV21 and DIV28 such that at DIV28 the difference between BACHD and WT was significant (WT:87.15±4.07; BACHD:76.4±5.69; p=0.035) (Fig. 5F and **Table 2**). Taken together the data suggest both the presynaptic and postsynaptic compartments, especially the latter, contributed to the reduction in synapses in BACHD cultures.

**Table 2.** Analysis of synaptic formation in cultured neurons of WT, BACHD, ΔN17 BACHD (II)

|  | DIV14 |  |  | DIV21 |  |  | DIV28 |  |  |
| --- | --- | --- | --- | --- | --- | --- | --- | --- | --- |
| | WT | BACHD | ΔN17BACHD | WT | BACHD | ΔN17BACHD | WT | BACHD | ΔN17BACHD |
| PSD95 (pixel) | 8.48±0.383 | 10.27±0.677<sup>ns</sup> | 9.93±1.16<sup>ns</sup> | 16.56±1.22 | 13.78±1.6<sup>ns</sup> | 20.59±2.12<sup>ns</sup> | 19.9±3.61 | 10.11±0.59<sup>**</sup> | 20.75±1.83<sup>ns</sup> |
| Synapsin I (pixel) | 7.14±0.33 | 9.54±0.45<sup>**</sup> | 7.10±0.58<sup>ns</sup> | 5.85±0.32 | 4.90±0.41<sup>ns</sup> | 8.47±0.66<sup>**</sup> | 6.64±0.94 | 6.95±0.6<sup>ns</sup> | 9.3±0.6<sup>ns</sup> |
| #synapses/100μm | 73.68±10.12 | 63.79±3.51<sup>ns</sup> | 67.11±6.58<sup>ns</sup> | 92.82±20.08 | 90.46±7.94<sup>ns</sup> | 93.93±7.87<sup>ns</sup> | 87.15±4.07 | 76.4±5.69<sup>*</sup> | 100.9±5.36<sup>**</sup> |

Significance test: One Way Anova (Dunnett's post test). ns= non-significance, $p > 0.05$ ; \* $p < 0.05$ ; \*\* $p < 0.01$

In addition to synapse number, ΔN17-BACHD neurons differed from WT in other synaptic parameters (Fig 5 and **Table 1**). Consistent with the increase in synapse number there was an increase in synapse density at DIV 28 (WT: 87.15±4.07; ΔN17-BACHD: 100.9±5.36, p=0.043) (Fig. 5D, **Table 2**). There were marked differences in PSD95/Synapsin 1 relative to WT; (ΔN17-BACHD vs WT): a decrease at DIV14: 0.119±0.010 vs 0.238±0.027 (Fig. 1E) (p<0.0001); no change at DIV21: 0.124±0.011 vs 0.125±0.010 (Fig. 2 E) (P>0.05); an increase at DIV28: 0.296 ± 0.024 vs 0.107 ± 0.019; (Fig. 3 E) (p<0.0001) and at DIV 35 (Fig. 4E) (0.260 ± 0.016 vs 0.163±0.019; p<0.0001) (Fig. 5B). There were also increases in Synapsin 1/PSD95 during the same period; the changes largely mirrored those in WT cultures (Fig. 5C, **Table 1**) and only at DIV21 was the difference significant (WT: 0.667±0.019 vs ΔN17-BACHD:0.78±0.018, p<0.01). The sizes of PSD95 puncta did not differ from WT (Fig. 5D, **Table 2**); Synapsin 1 puncta were larger than WT at DIV21 (WT:5.85±0.32; ΔN17-BACHD8.47±0.66, p<0.01) (Fig. 5E, **Table 2**). Synapsin I puncta remained larger than WT at DIV28 albeit not significantly (9.3±0.6 vs 6.64±0.94, p=0.06). These data suggest that delta N17 BACHD neurons differed from WT in both the pre- and postsynaptic compartments with changes in the latter contributing more significantly to increases in synapse number and density. As such, the data suggest an increase in synapse formation, maintenance or both in ΔN17-BACHD cultures relative to the WT.

### Age-related changes in synaptic proteins in WT and BACHD neurons

The significant age-related reduction of synapses in BACHD raised the possibility that synaptic proteins were reduced in BACHD neurons. To test this possibility, we carried out Western blotting analysis of primary culture neuronal cultures. E18 cortical neurons from WT and BACHD mice were cultured as described above. Neuronal cultures were harvested and lysates assayed by immunoblotting with antibodies specific for synaptic proteins involved in SNARE complex formation (synaptobrevin, SNAP25), calcium sensing (synaptotagmin), synaptic vesicles (synaptophysin) and the post-synapse (PSD95) (Fig. S1). We chose to examine DIV21 andDIV28 cultures to examine synaptic proteins at times shown to demonstrate significant differences in synapse number between the mutant and WT neurons. Remarkably, at DIV21, BACHD neurons showed a statistically significant increases in all proteins examined (Fig. S1). However, at DIV28 the levels of all these proteins were significantly decreased (Fig. S1). The latter is consistent with the decrease in synapse numbers at DIV28, but exceeds in magnitude the reduction in synapse number. These data are evidence for age-related changes in synaptic proteins in BACHD cultures but do not demonstrate a direct relationship between synaptic protein levels and synapse number in BACHD neurons.

### BDNF treatment prevented synaptic deficits in BACHD cortical neurons

BDNF signaling plays an important role in mediating activity-dependent synaptic plasticity, including synaptogenesis and synaptic function (Park and Poo, 2013). Previously, we showed that anterograde axonal transport of BDNF was reduced in BACHD cortical axons and provided evidence that this deficit was linked to decreased BDNF release to support the trophic status of striatal neurons(Zhao et al., 2016). To test directly for decreased BDNF release from BACHD cortical axons as possibly contributing to synapse loss we collected conditioned media from WT and BACHD cortical neurons at DIV14 and DIV 21 and quantitated BDNF by ELISA, as described in the Materials and Methods. To normalize secreted BDNF to total cell protein we harvested protein lysates from cultures and measured protein content. At DIV 14 the level of released BDNF in BACHD cultures did not differ significantly from WT neurons (WT:3.863±0.5288 pg BDNF/mg protein; BACHD: 3.542±0.4215 pg/mg protein; p=0.6627) (Fig. S2). However, at DIV21 BDNF released in BACHD cultures was significantly lower than in WT neurons (WT: 4.433±0.3310 pg BDNF/mg protein; BACHD: 3.233±0.1389 pg BDNF/mg proteins; p=0.0189) (Fig. S2). Though the differences between WT and BACHD were significant, the changes between DIV14 and DIV21 were not significant for either WT or BACHD neurons (DIV14 vs DIV21; WT: p=0.3877; BACHD: p=0.5123). We conclude that in comparison to WT cultures in BACHD cultures BDNF release is decreased with increased age.

In earlier studies we demonstrated that adding BDNF to BACHD striatal neurons reversed atrophy (Zhao et al., 2016). To ask treating with exogenous BDNF would prevent synaptic changes, we treated WT and BACHD cortical neurons with BDNF (100 ng/ml) for either 7 (DIV 14 to DIV 21) or 14 days (DIV 7 to DIV 21) and immunostained as above for PSD95 and Synapsin 1. The findings were compared to those in untreated cultures. BDNF treatment for either 7 (Fig. 6) or 14 days (Fig. 7) resulted in a significant increase in synapse number in both WT and BACHD neurons (Fig. 6C, 7C; Table 3). With BDNF treatment for 7 days (Fig. 6), the colocalization coefficient was 0.675 ± 0.011 in BACHD neurons, a value comparable to that (0.651±0.008) (p=0.087) in WT neurons (Fig. 6C, Table 3). With BDNF treatment for 14 days (Fig. 7), the colocalization coefficient in BACHD neurons was increased further (0.705 ± 0.009) and was significantly higher than that in WT cultures (0.665 ± 0.010) (p=0.005) in WT neurons (Fig. 7C, Table 3).

**Figure 6.**
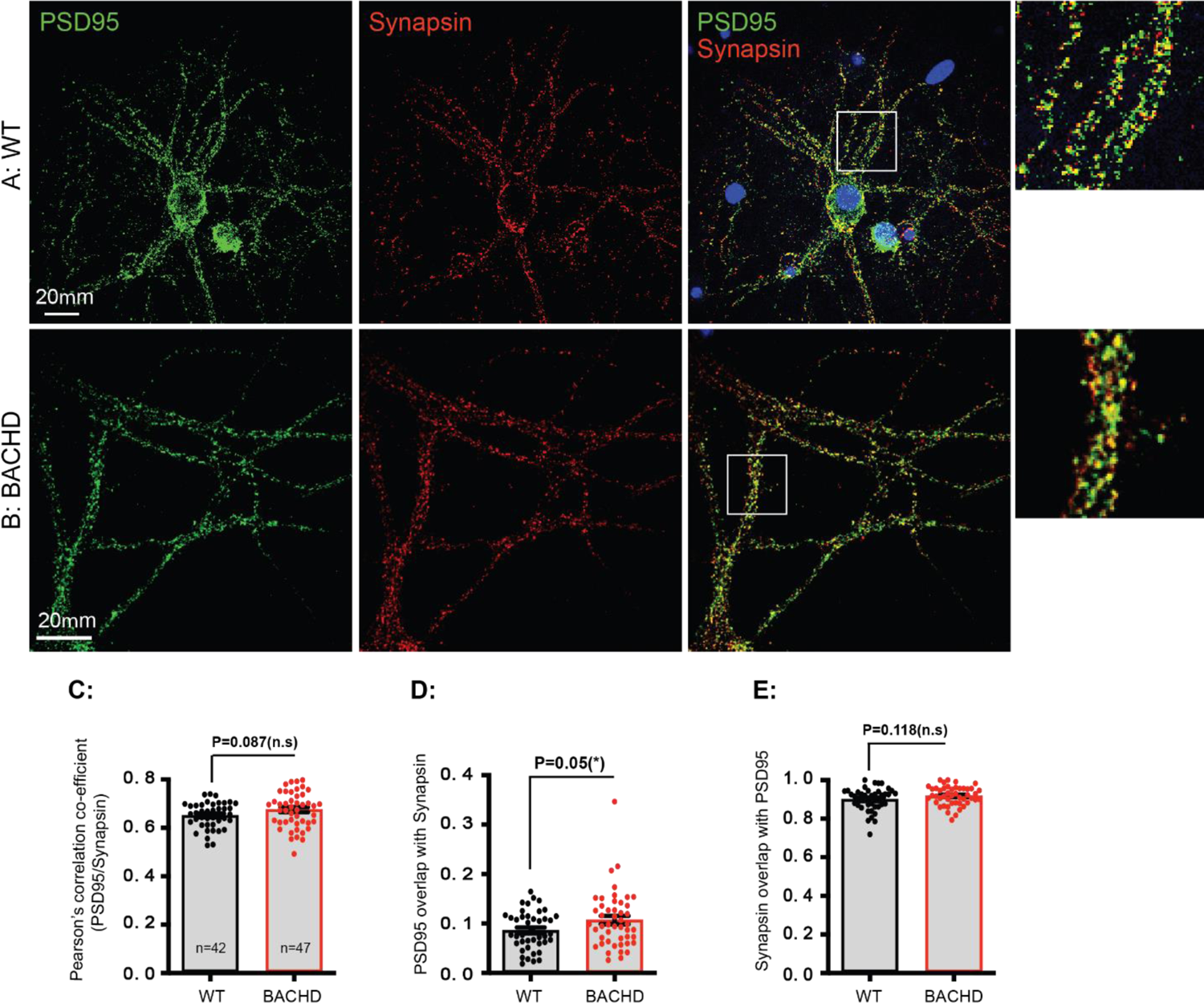
Rescuing effect of synaptic deficits in BACHD neurons by a 7-day treatment with BDNF. E18 cortical neurons from WT and BACHD were dissected, cultured as in Figure 1-4. Starting at DIV14, 100ng/ml BDNF was added to the maintenance media and the media were replaced every other day until DIV21. Neurons were then fixed, immune-stained and quantitated for Pearson’s colocalization co-efficient as for Figure 1-4. (A) (B) show representative images of PSD95 (green) and Synapsin I (red) staining respectively in BACHD cultures with vehicle or BDNF treatment. Regions of interest marked by white boxes are magnified and shown on the right. (C) Comparison of post- and presynaptic marker colocalization using Pearson’s Colocalization Coefficients (PCC). (D) Analysis of PSD95 signals that overlapped with Synapsin I. (E) Analysis of Synapsin I signals that overlapped with PSD95. Results are shown as mean ± SEM. The numbers of images were analyzed for n=42 (WT), n=47 (BACHD). The data represent at least 4-5 independent cultures. Significance analysis was carried out using Prism. Statistical significances were calculated by unpaired Student’s t test. n.s.= non significance. All p values are shown in the graphs.

**Figure 7.**
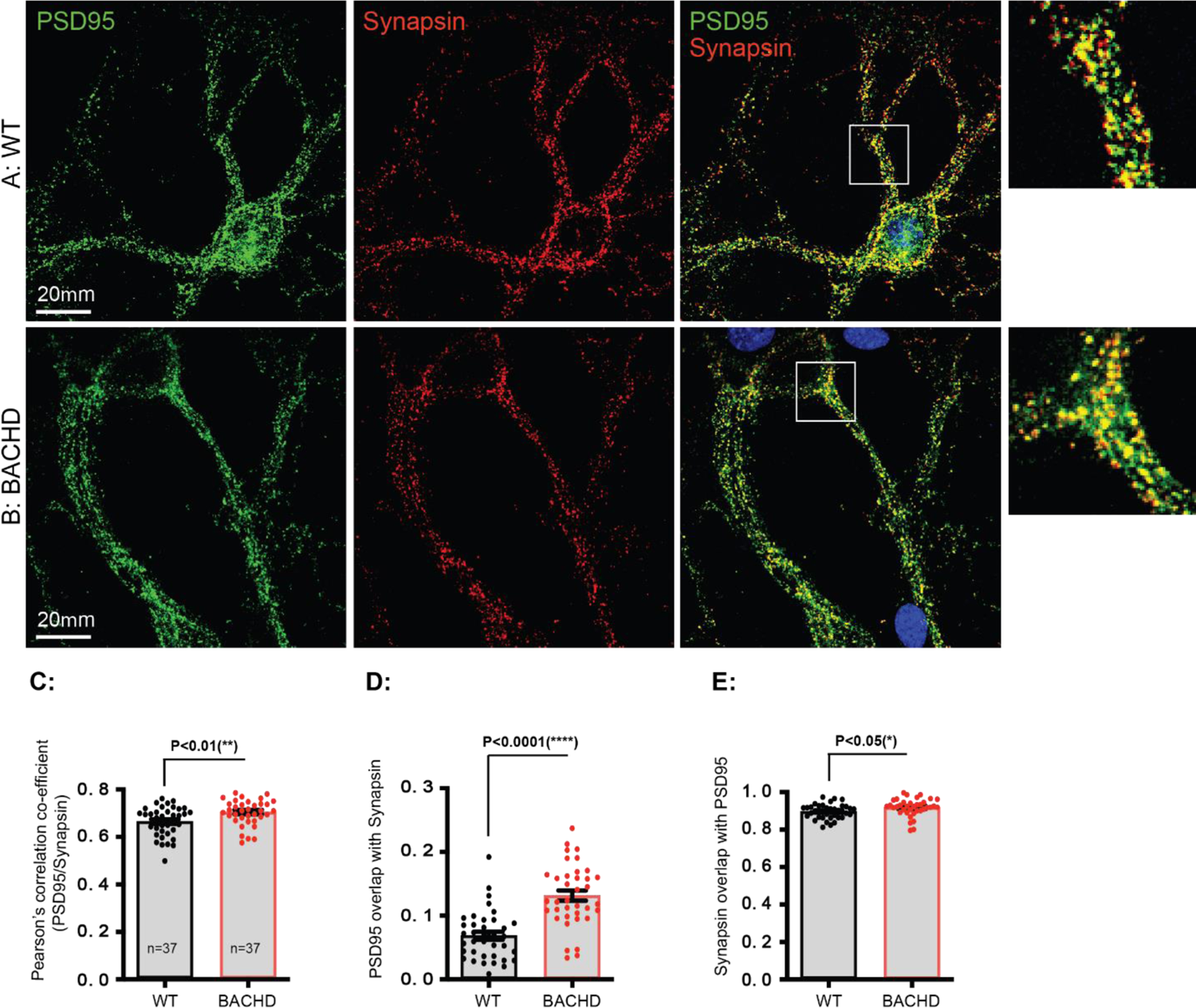
Rescuing effect of synaptic deficits in BACHD neurons by a 14-day treatment with BDNF. E18 cortical neurons from WT and BACHD were dissected, cultured as in Figure 1-4. Starting at DIV7, 100ng/ml BDNF was added to the maintenance media and the media were replaced every other day until DIV21. Neurons were then fixed, immune-stained and quantitated for Pearson’s colocalization co-efficient as for Figure 2-5. (A) (B) show representative images of PSD95 (green) and Synapsin I (red) staining respectively in BACHD cultures with vehicle or BDNF treatment. Regions of interest marked by white boxes are magnified and shown on the right. (C) Comparison of post- and presynaptic marker colocalization using Pearson’s Colocalization Coefficients (PCC). (D) Analysis of PSD95 signals that overlapped with Synapsin I. (E) Analysis of Synapsin I signals that overlapped with PSD95. Results are shown as mean ± SEM. The numbers of images were analyzed: n=37 (WT), n=37 (BACHD). The data represent at least 4-5 independent cultures. Significance analysis was carried out using Prism. Statistical significances were calculated by unpaired Student’s t test. n.s.= non significance. All p values are shown in the graphs.

**Table 3.** Analysis of BDNF effects on synaptic formation in WT, BACHD

|  | WT |  |  | BACHD |  |  |
| --- | --- | --- | --- | --- | --- | --- |
|  | Vehicle | BDNF 7 d | BDNF 14 d | Vehicle | BDNF 7d | BDNF 14d |
| PCC | 0.565±0.010 | 0.651±0.008*** | 0.665±0.010*** | 0.422±0.019 | 0.675±0.011*** | 0.705±0.009*** |
| PSD95/Synapsin I | 0.125±0.010 | 0.086±0.006** | 0.069±0.006*** | 0.018±0.003 | 0.107 ± 0.008*** | 0.132 ± 0.008*** |
| Synapsin I/PSD95 | 0.667±0.019 | 0.900 ± 0.009*** | 0.896 ± 0.006*** | 0.535±0.041 | 0.918 ± 0.007*** | 0.918 ± 0.008*** |

Significance test: One Way ANOVA (Dunnett's post test). ns= non-significance, $p>0.05$ ; \* $p<0.05$ ; \*\* $p<0.01$ ; \*\*\* $p<0.001$ );

BDNF treatment also impacted other synaptic parameters. The percent of PSD95/Synapsin I increased following BDNF treatment (Fig. 6D, 7D; Table 3). In BACHD neurons treated for 7 days, this value (0.107 ± 0.008) was significantly greater than in WT neurons (0.086 ± 0.006) (p=0.05) (Fig. 6D, Table 3). Treatment for 14 days resulted in a value of 0.132 ± 0.008 in BACHD neurons, a result markedly greater than in WT neurons (0.069 ± 0.006) (p<0.0001) (Fig. 7D, Table 3). The differences relative to WT cultures were correlated with decreases in the PSD95/Synapsin 1 following BDNF treatment of WT cultures; the reduction after 7 days of treatment was less than after 14 days (WT BDNF treated vs untreated for 7 days: 0.0861±0.006 versus 0.1249±0.0101; p=0.0367); (WT BDNF treated versus untreated for 14 days: 0.069±0.006versus 0.125±0.010, p=0.004). BDNF treated BACHD neurons also increased Synapsin I/PSD95 after both 7 and 14 days relative to untreated cultures (untreated: 0.535±0.004; treated 7 days: 0.918 ± 0.007, p<0.0001; treated 14 days: 0.918 ± 0.008, p<0.0001) (Fig. 6E, 7E, Table 3), resulting in values in BACHD neurons that were equal to or greater than in WT neurons (WT untreated: 0.667±0.019; WT treated 7d, 0.900 ± 0.009, p=0.118; WT treated 14 d; 0.896 ± 0.006, p=0.031) (Fig. 6E, 7E, Table 3). WT neurons thus also responded to BDNF with respect to the percent Synapsin1/PSD 95.

In summary, BACHD neurons responded to BDNF treatment with significant increases in synapse number (i.e. increases in PCC) PSD95/Synapsin 1 and Synapsin 1/PSD95. The increases resulted in values equal to or greater than in WT controls (Table 3) through effects that appear to have involved both the presynaptic and postsynaptic compartments. Remarkably, and consistent with earlier studies on its trophic effects on striatal neurons (Zhao et al., 2016), BACHD neurons were equally or more responsive to BDNF than WT neurons.

### Synaptic deficits in BACHD cortical neurons were partially prevented by ApiCCT1

The chaperonin TRiC is a modulator of mHTT aggregation and toxicity(Shen et al., 2016). We demonstrated that expression of individual TRiC subunit(s) CCT3, CCT5 or exogenous addition of the apical domain of CCT1 (ApiCCT1) rescued BDNF transport in BACHD cortical neurons in cortico-striatal cultures (Zhao et al., 2016). The effect of ApiCCT1 on striatal neuron atrophy was correlated with increased anterograde axonal transport of BDNF and BDNF release(Zhao et al., 2016). To ask if, like BDNF, ApiCCT1 treatment would rescue synaptic deficits, we treated BACHD cortical neurons at DIV14 with either 0.1 μM ApiCCT1 or the vehicle control (Material and Methods). Synaptic staining and quantitation were carried out at DIV 21. As compared to vehicle, treatment with 0.1 μM ApiCCT1 significantly increased synapse number as reflected in PCC values (vehicle vs ApiCCT1: 0.490 ± 0.011 vs 0.563 ± 0.009, p<0.0001) (Fig. 8A). The effect of ApiCCT1 was correlated with a significant increase in the percent of PSD95/Synapsin I (0.312 ± 0.018 vs 0.523 ± 0.023, p<0.0001) (Fig. 8B); the percent in Synapsin I/PSD95 was modestly but significantly reduced (0.466 ± 0.025 vs 0.401 ± 0.017, p=0.048) (Fig. 8C). ApiCCT1 had no effect on puncta size for either PSD95 (11.14 ± 0.93 vs 10.74 ± 0.62, p=0.74) (Fig. 8D) or Synapsin I (8.73 ± 0.43 vs 8.63 ± 0.36, p=0.85) (Fig. 8E). (Fig. 8F) We conclude that ApiCCT1 treatment partially prevented the deficits in synapses in BACHD cortical neurons.

**Figure 8.**
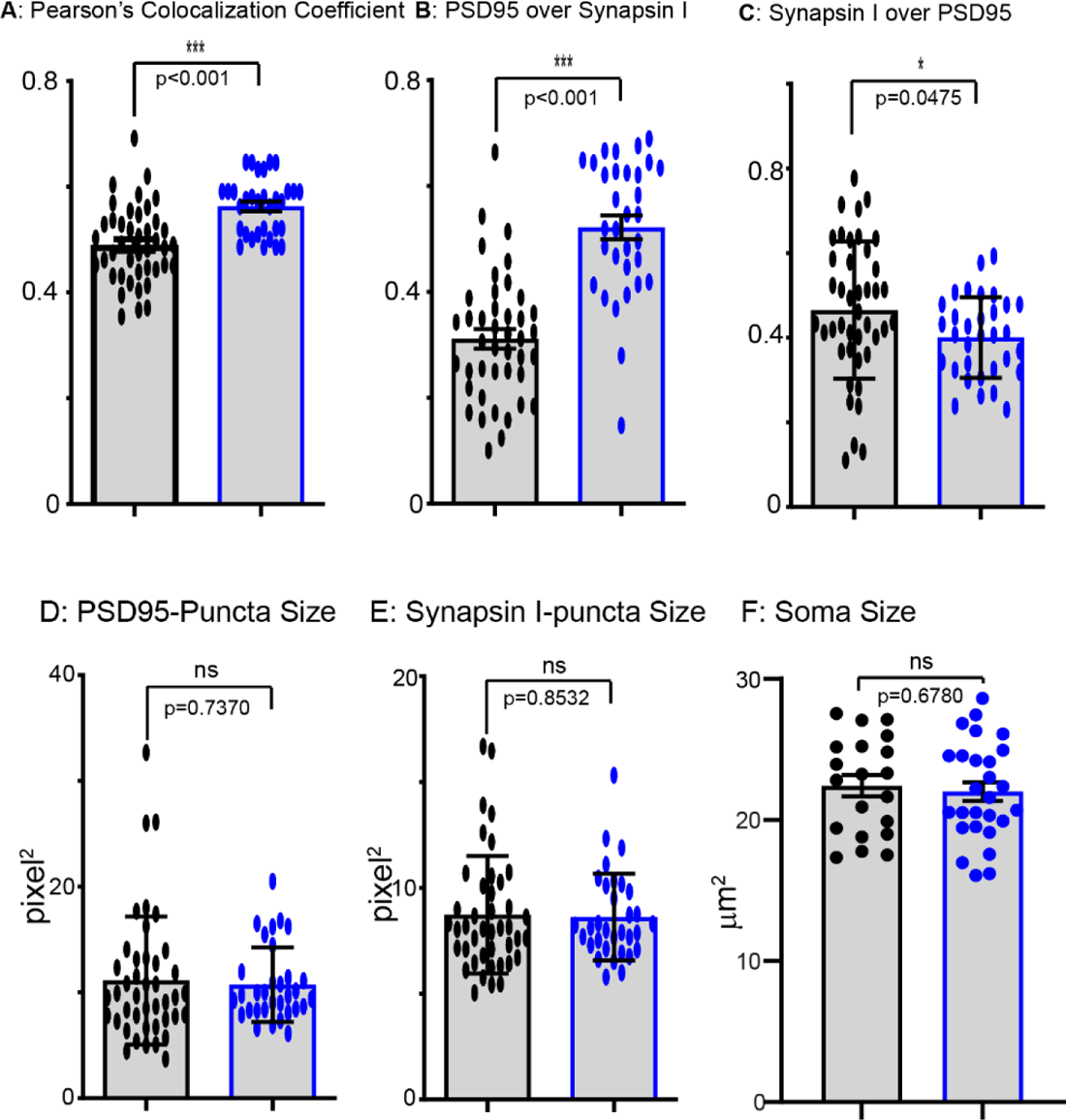
Rescuing effect of synaptic deficits in BACHD neurons by ApiCCT1. E18 cortical neurons from WT and BACHD were dissected, cultured as in Figure 1-4. Starting at DIV14, 0.1µM ApiCCT1 was added to the maintenance media and the media were replaced every other day until DIV21. Neurons were then fixed, immune-stained and quantitated for Pearson’s colocalization co-efficient as for Figure 1-4. (A) Comparison of post- and presynaptic marker colocalization using Pearson’s Colocalization Coefficients (PCC) in BACHD cultures with vehicle or ApiCCT1 treatment. (B) Analysis of PSD95 signals that overlapped with Synapsin I. (C) Analysis of Synapsin I signals that overlapped with PSD95. (C) Measurements of PSD95 puncta sizes. (E) Measurements of Synapsin I puncta sizes. (F) Analysis of soma sizes of cortical neurons in BACHD cultures treated with vehicle or ApiCCT1. Results are shown as mean ± SEM. The numbers of images were analyzed: n=42 (WT), n=32 (BACHD). The data represent at least 4-5 independent cultures. Significance analysis was carried out using Prism. Statistical significances were calculated by unpaired Student’s t test. n.s.= non significance. All p values are shown in the graphs.

### BACHD cortical neurons showed significant functional deficits at DIV28

We next asked if BACHD cortical neurons were deficient in synaptic function. To this end, we used multielectrode arrays (MEA)(Cotterill et al., 2016). The array contains a grid of tightly spaced electrodes embedded in the culture surface to record neuronal activity. When neurons fire action potentials, the extracellular voltage is measured by the electrodes on a microsecond timescale(Cotterill et al., 2016). MEA is well suited for measuring neuronal network activity and for sampling events at many locations across the culture to record initiation, propagation and synchronization of neural activity.

E18 cortical neurons from WT and BACHD were cultured in CytoView MEA 24-well plates pre-coated with poly-D-lysine (Axion Biosystems) at a density of 100,000 cells per well. At DIV14, neuronal activity and key features of neural network behaviors such as activity, synchrony, and network oscillations were recorded on the Maestro Edge (Axion) for 10 minutes. The data were batch processed using AxIS Navigator v2.0.4.21. We quantitated the following metrics of neuronal activities: 1) weighted mean firing rate (wMFR: average spikes per sec per active electrode) to estimate the overall population excitability and connectivity; 2) Interspike intervals (ISIs) coefficient of variation (ISI CoV) to measure irregularity of spike trains; 3) synchrony index to measure the degree to which activity is synchronized; the number of spikes per burst; 4) total number of bursts; 5) burst frequency; 6) numbers of spikes/burst; 7) inter-burst intervals (IBIs) to measure the length of quiescent periods between bursts; 8) percentage of network bursts – i.e. the number of spikes in network bursts divided by the total number of spikes, multiplied by 100; 9) numbers of spikes/network burst; 10) the mean ISI within a burst to detect changes in burst patterns; 11) network bursts, to measure periodic and synchronized activity of cultured neurons; and 12) network inter-burst intervals (IBIs) coefficient of variation (IBI CoV). As shown in Fig. 9, BACHD neurons at DIV14 showed a difference from WT in only one parameter (Fig. 9A-L). BACHD neurons differed in the network IBI CoV with a significant reduction denoting more regular network bursting (Fig. 9 L). These findings provide function data complementing those for synapse number and synapse density at DIV 14. They are evidence that BACHD cortical neurons form functional synapses with properties very similar to WT cultures.

**Figure 9.**
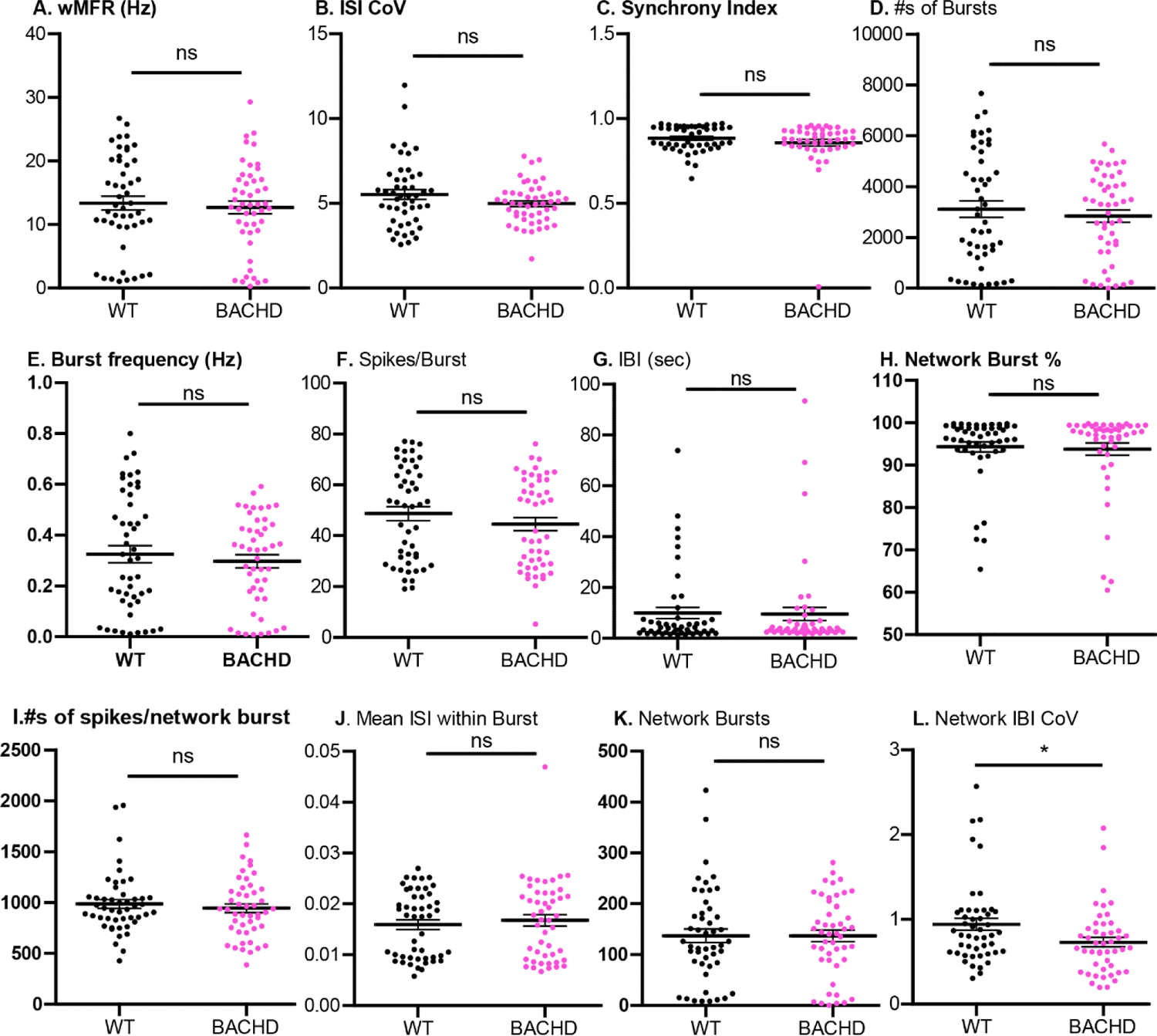
BACHD cortical neuronal activity shows no significant deficits at DIV14 E18 cortical neurons from WT and BACHD were plated on Poly-D-Lysine coated (100 µg/ml – Sigma Aldrich) CytoView MEA 24 well plates (Axion Biosystems) at a density of 100,000 cells per well. At DIV14, neuronal activity and key features of neural network behavior as functional endpoints - activity, synchrony, and network oscillations were recorded on the Maestro Edge (Axion) for 10 minutes. The data were batch processed using AxIS Navigator v2.0.4.21. E18 BACHD cortical neurons showed no difference from WT in weighted mean firing rate (A), ISI (inter spike interval) coefficient of variation (B), the number of spikes/burst (C), mean ISI within burst, numbers of bursts (E), inter-burst interval ((F), network bursts (G), synchrony index (H). However, BACHD cortical neurons showed a significant reduction in the network inter burst interval (IBI) coefficient of variation (I). Each data point represents one well of data. Analysis by two-way ANOVA. * indicates p<0.05, ns=not significant.

At DIV28, several changes in synapse function characterized BACHD cultures with respect to WT cultures (Fig. 10). BACHD neurons showed significant changes: 1) decreased ISI CoV (Fig. 10B); 2) decreased synchrony index (Fig. 10C); 3) increased IBI (Fig. 10G); and decreased network burst percentage (Fig. 10H). There were no significant deficits in the following metrics: wMFR (Fig 10A); numbers of burst (Fig. 10D); burst frequency (Fig. 10E); spikes/burst (Fig. 10F); numbers of spikes/network burst (Fig 10 I); mean ISI within burst (Fig. 10J), network bursts (Fig. 10K) or network IBI coefficient of variation (Fig. 10L). The changes point to dysregulation of synaptic activity with differences in timing of spikes and bursts, the percentage of bursts and network synchrony. Nevertheless, a number of functional measures were preserved. Table 4 compares the findings at DIV 28 to those at DIV 14 for both WT and BACHD.

**Figure 10.**
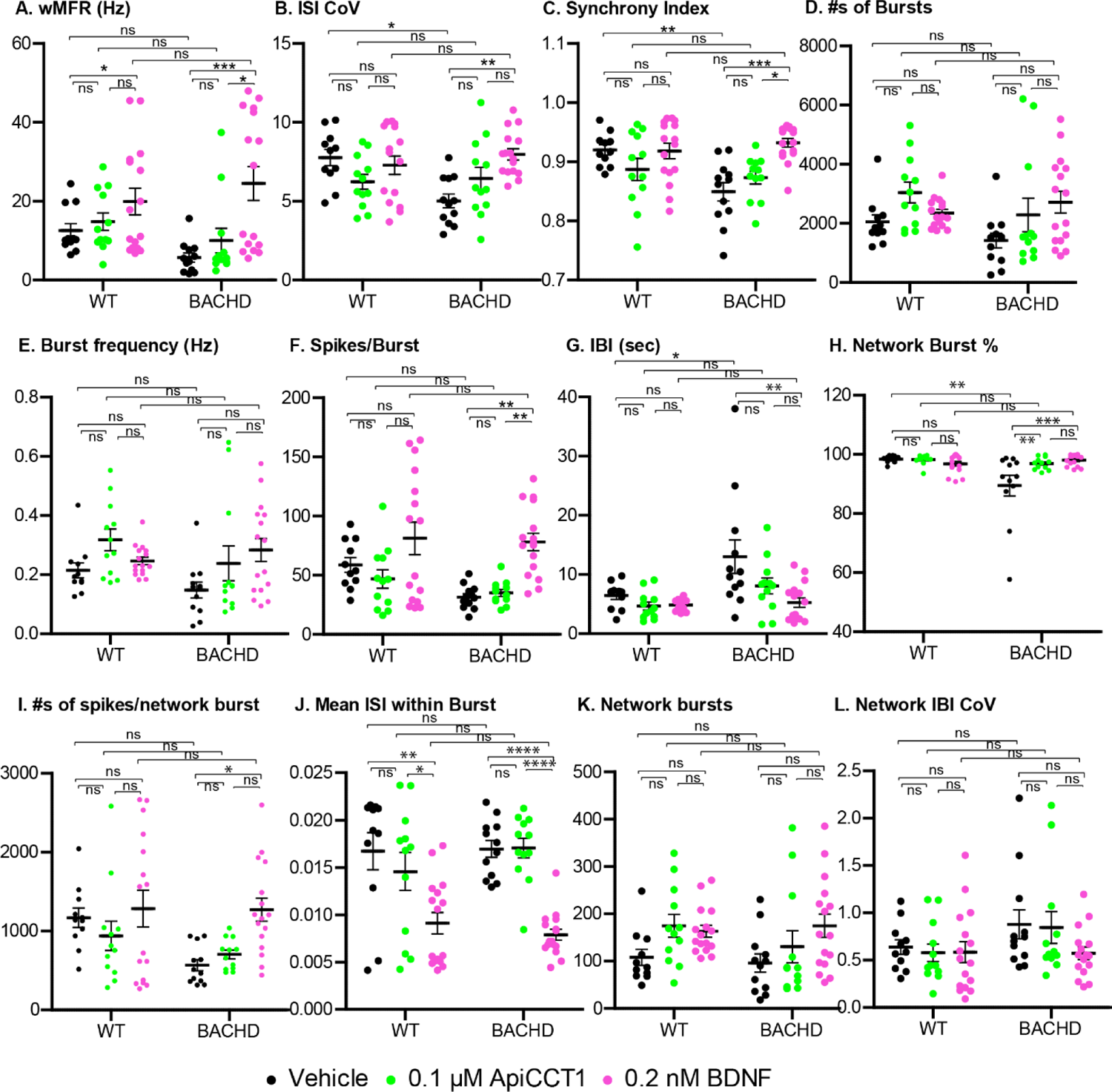
BACHD cortical neuronal activity shows significant deficits at DIV28. After completion of recording at DIV14, WT and BACHD neuronal cultures were treated with BDNF (50 ng/ml), ApiCCT1 (0.1µM) or Vehicle. Media were changed every 48 hrs and a final recording was performed as above at DIV 28. Data were collected and quantitated: weighted mean firing rate (A), ISI (inter spike interval) coefficient of variation (B), the number of spikes/burst (C), mean ISI within burst, numbers of bursts (E), inter-burst interval ((F), network bursts (G), synchrony index (H), network inter burst interval (IBI) coefficient of variation (I). Each data point represents one well of data. Analysis by two-way ANOVA. *: p<0.05, **: p<0.01, ***: p<0.001, ****: p<0.0001, ns=not significant.

**Table 4.** Comparison of Neuronal Activities

| MEA metrics | WT: DIV14 vs DIV28 |  |  | BACHD: DIV14 vs DIV28 |  |  | WT DIV28 vs BACHD DIV28 |  |  |
| --- | --- | --- | --- | --- | --- | --- | --- | --- | --- |
|  | DIV14 | DIV28 | p value | DIV14 | DIV28 | p value | WT | BACHD | p value |
| A. wMFR (Hz) | 13.38±1.08 | 12.55±1.79 | 0.3354 ns | 12.69±1.01 | 5.72±1.15 | 0.0027 ** | 12.55±1.79 | 5.72±1.15 | 0.0015 ** |
| B. ISI CoV | 5.51±0.29 | 7.77±0.52 | 0.0009 *** | 4.98±0.17 | 5.02±0.44 | 0.8898 ns | 7.77±0.52 | 5.02±0.44 | 0.0012 ** |
| C. Synchrony Index | 0.88±0.01 | 0.92±0.01 | 0.3552 ns | 0.86±0.02 | 0.85±0.02 | 0.1574 ns | 0.92±0.01 | 0.85±0.02 | 0.0012 ** |
| D. #s of Bursts | 3112±324 | 2046±242 | 0.2631 ns | 2839±244 | 1425±254 | 0.0117 * | 2046±242 | 1425±254 | 0.0337 * |
| E. Burst Frequency (Hz) | 0.33±0.03 | 0.25±0.03 | 0.2888 ns | 0.30±0.03 | 0.15±0.03 | 0.0117 * | 0.25±0.03 | 0.15±0.03 | 0.0337 * |
| F. Spikes/Burst | 48.70±2.70 | 58.73±6.11 | 0.1582 ns | 44.55±2.57 | 31.59±2.90 | 0.026 * | 58.73±6.11 | 31.59±2.90 | 0.0012 ** |
| G. IBI (sec) | 9.95±2.18 | 6.44±0.67 | 0.1001 ns | 9.52±2.57 | 13.03±2.83 | 0.0025 ** | 6.44±0.67 | 13.03±2.83 | 0.0289 ns |
| H. Network Burst% | 94.31±1.17 | 98.35±0.36 | 0.0886 ns | 93.76±1.44 | 89.42±3.50 | 0.0375 * | 98.35±0.36 | 89.42±3.50 | 0.0028 ns |
| I. Spikes/Network Burst | 986±43 | 1169±121 | 0.0749 ns | 944±43 | 566±70 | 0.0004 *** | 1169±121 | 566±70 | 0.0005 *** |
| J. Mean ISI w/n Burst | 0.016±0.001 | 0.017±0.002 | 0.8916 ns | 0.017±0.001 | 0.017±0.001 | 0.9484 ns | 0.017±0.002 | 0.017±0.001 | 0.5181 ns |
| K. Network Bursts | 137±13 | 108±17 | 0.2351 ns | 137±11 | 96±19 | 0.0873 ns | 108±17 | 96±19 | 0.5795 ns |
| L. Network IBI CoV | 0.94±0.07 | 0.64±0.08 | 0.0286 * | 0.73±0.06 | 0.88±0.15 | 0.5405 ns | 0.64±0.08 | 0.88±0.15 | 0.2549 ns |

Significance test: nonparametric t-test (Mann-Whitney) ns= non-significance, $p>0.05$ ; \* $p<0.05$ ; \*\* $p<0.01$ ; \*\*\* $p<0.001$ )

Given that BDNF and ApiCCT1 treatment of BACHD cultures positively impacted synapse number and structure, we asked if these treatments would prevent changes in synaptic activity. For these studies, BDNF treatment was initiated at DIV14 and cultures were examined at DIV28. BDNF treatment of BACHD neurons showed a robust effect in normalizing deficits. Fig S3 shows representative screen shots of firing patterns of WT, BACHD neurons treated with either vehicle or BDNF (Fig S3). In vehicle treated cultures, the amplitudes of BACHD activity were reduced and the pattern was irregular as compared to WT neurons (Fig. S1, B vs A). These effects were completely largely ameliorated with BDNF treatment (Fig. S1, D vs B). There was no obvious change in activity pattern for WT neurons in response to BDNF (Fig. S1, C vs A). Using Two Way ANOVA to explore synaptic activity for significant and noteworthy effects of BDNF treatment were detected for measures that distinguished BACHD from WT neurons (Fig. 10). Thus, BDNF treatment: 1) increased ISI coefficient of variation (Fig. 10B); 2) increased synchrony index (Fig. 10C); 3) decreased IBI (Fig. 10G); and increased network burst percentages (Fig. 10H). BDNF also impacted measures that did not distinguish BACHD and WT neurons. BDNF treatment of BACHD neurons significantly improved wMFR (Fig. 10A); increased spikes/burst (Fig. 10F); and increased numbers of spikes/network burst (Fig. 10I). In addition, BDNF markedly reduced mean ISI within bursts to values lower than in vehicle-treated WT cultures (Fig. 10J). BDNF treatment of WT cultures showed fewer effects; it significantly increased wMFR (Fig. 10A) and as for BACHD cultures decreased mean ISI within bursts (Fig. 10J). BDNF treatment thus prevented BACHD-mediated synaptic changes and increased additional parameters reflecting synaptic function.

To examine further the effects of BDNF treatment, and to specifically address the effects within genotype, we used One-Way ANOVA to examine WT and BACHD synaptic activity (Table 5). Among the 12 MEA metrics measured, BDNF treatment of WT neurons induced a significant change only in mean ISI within burst (Vehicle: 0.017±0.002 vs BDNF: 0.009±0.001, p =0.0045). However, treatment of BACHD neurons significantly impacted many MEA metrics (Table 5): 1) mWFR (Vehicle: 5.72±1.15 vs BDNF: 22.97±4.29, p=0.0008); 2) ISI CoV (Vehicle: 5.02±0.44 vs BDNF 7.99±0.38, p=0.003); 3) Synchrony index (Vehicle: 0.85±0.02 vs BDNF: 0.93±0.01, p<0.0001); 4) spikes/burst (Vehicle: 31.59±2.90 vs BDNF: 78.2±7.39, p<0.0001); 5) IBI (Vehicle: 13.03±2.83 vs BDNF: 5.25±0.80, p=0.0046); 6) Network burst % (Vehicle: 89.42±3.50 vs BDNF 98.04±0.47, p=0.04); 7) Spikes/network burst (Vehicle: 566±70 vs BDNF:1272±147, p=0.0001); 8) Mean ISI within burst (Vehicle: 0.017±0.001 vs BDNF: 0.0079±0.001, p<0.0001). These data are evidence that BACHD neurons are robustly responsive to BDNF treatment with respect to synaptic function.

**Table 5.**
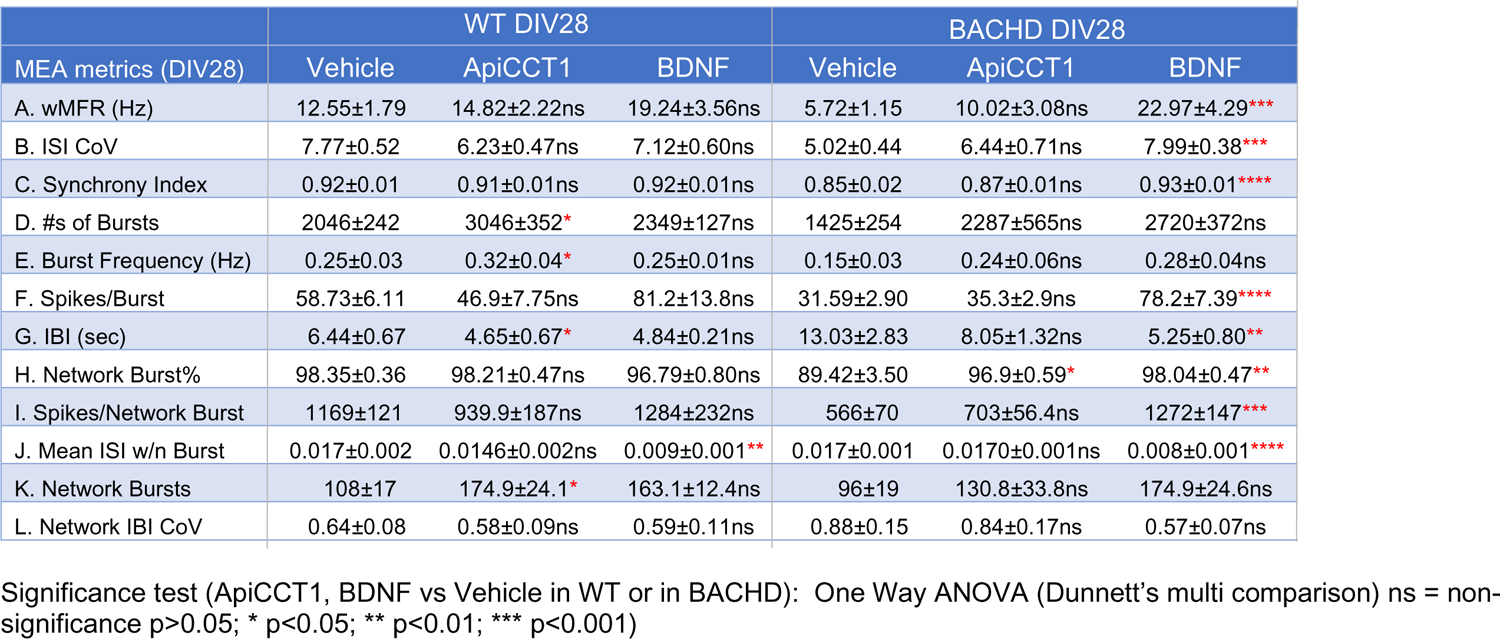
Effects of ApiCCT1, BDNF on neuronal activities

| MEA metrics (DIV28) | WT DIV28 |  |  | BACHD DIV28 |  |  |
| --- | --- | --- | --- | --- | --- | --- |
|  | Vehicle | ApiCCT1 | BDNF | Vehicle | ApiCCT1 | BDNF |
| A. wMFR (Hz) | 12.55±1.79 | 14.82±2.22ns | 19.24±3.56ns | 5.72±1.15 | 10.02±3.08ns | 22.97±4.29*** |
| B. ISI CoV | 7.77±0.52 | 6.23±0.47ns | 7.12±0.60ns | 5.02±0.44 | 6.44±0.71ns | 7.99±0.38*** |
| C. Synchrony Index | 0.92±0.01 | 0.91±0.01ns | 0.92±0.01ns | 0.85±0.02 | 0.87±0.01ns | 0.93±0.01**** |
| D. #s of Bursts | 2046±242 | 3046±352* | 2349±127ns | 1425±254 | 2287±565ns | 2720±372ns |
| E. Burst Frequency (Hz) | 0.25±0.03 | 0.32±0.04* | 0.25±0.01ns | 0.15±0.03 | 0.24±0.06ns | 0.28±0.04ns |
| F. Spikes/Burst | 58.73±6.11 | 46.9±7.75ns | 81.2±13.8ns | 31.59±2.90 | 35.3±2.9ns | 78.2±7.39**** |
| G. IBI (sec) | 6.44±0.67 | 4.65±0.67* | 4.84±0.21ns | 13.03±2.83 | 8.05±1.32ns | 5.25±0.80** |
| H. Network Burst% | 98.35±0.36 | 98.21±0.47ns | 96.79±0.80ns | 89.42±3.50 | 96.9±0.59* | 98.04±0.47** |
| I. Spikes/Network Burst | 1169±121 | 939.9±187ns | 1284±232ns | 566±70 | 703±56.4ns | 1272±147*** |
| J. Mean ISI w/n Burst | 0.017±0.002 | 0.0146±0.002ns | 0.009±0.001** | 0.017±0.001 | 0.0170±0.001ns | 0.008±0.001**** |
| K. Network Bursts | 108±17 | 174.9±24.1* | 163.1±12.4ns | 96±19 | 130.8±33.8ns | 174.9±24.6ns |
| L. Network IBI CoV | 0.64±0.08 | 0.58±0.09ns | 0.59±0.11ns | 0.88±0.15 | 0.84±0.17ns | 0.57±0.07ns |

Significance test (ApiCCT1, BDNF vs Vehicle in WT or in BACHD): One Way ANOVA (Dunnett's multi comparison) ns = non-significance $p>0.05$ ; \* $p<0.05$ ; \*\* $p<0.01$ ; \*\*\* $p<0.001$ )

To test the effects of ApiCCT1, we employed media containing either ApiCCT1 or vehicle. Media were first added on DIV 14 and were refreshed every other day until recording at DIV28. ApiCCT1 treatment of BACHD neurons significantly increased only network burst percentage (Fig 10H). While there was no apparent ApiCCT1 effect on WT cultures using Two-Way ANOVA (Fig. 10) analysis, using One-Way ANOVA to emphasize effects within a genotype (Table 5) showed that ApiCCT1 treatment of WT neurons induced significant changes in: 1) #s of bursts (Vehicle: 2046±242 vs ApiCCT1: 3046±352, p=0.0162) 2) Burst frequency (Vehicle:0.25±0.03 vs ApiCCT1:0.32±0.04, p=0.0173); 3) IBI (Vehicle: 6.44±0.67 vs ApiCCT1: 4.65±0.67, p=0.042); and 4) Network bursts (Vehicle:108±17 vs ApiCCT1:174.9±24.1, p=0.0302). These findings point not only to different treatment efficacies for BDNF and ApiCCT1 on BACHD and WT neurons but also for differential effects on synaptic function.

## Discussion

Synaptic homeostasis is critical for neuronal structure and function. Defects in synapse structure and function mark neurodegenerative disorders, including HD (Barry et al., 2022; Chen et al., 2019; Cummings et al., 2007; Kolodziejczyk et al., 2014; Lepeta et al., 2016; Milnerwood et al., 2006; Milnerwood and Raymond, 2007; Murphy et al., 2000; Nithianantharajah and Hannan, 2013; Schirinzi et al., 2016; Selkoe, 2002; Subramanian and Tremblay, 2021; Taoufik et al., 2018; Veldman and Yang, 2018); they contribute directly to neuronal circuit dysfunction and resulting changes in cognitive, behavioral and motor dysfunction(Chen et al., 2019; Cummings et al., 2007; Dallerac et al., 2011; Kavalali et al., 1999; Kolodziejczyk et al., 2014; Kovalenko et al., 2018; Lepeta et al., 2016; Li et al., 2003; Morton et al., 2001; Nithianantharajah and Hannan, 2013; Selkoe, 2002; Smith-Dijak et al., 2019; Subramanian and Tremblay, 2021). To address synaptic dysfunction and loss it is essential to define the molecular events responsible and discover methods by which to intercept pathogenesis. We employed primary cultures of cortical neurons from the BACHD and ΔN17-BACHD mouse models of HD to explore synaptic pathogenesis. Following an early period during which synapses apparently were formed and functioned normally, both models showed significant deviations from normal. BACHD neurons showed regressive changes in synapse number and function. Remarkably, BDNF treatment prevented most such changes. Indeed, BDNF treatment of BACHD neurons resulted in changes in synapse structure and function that in some cases exceeded those in vehicle-treated WT cultures. The TRiC-inspired reagent ApiCCT1 induced similar but generally less robust effects, especially with respect to synaptic function. Our findings demonstrate the ability to explore synapse maintenance and function in HD model systems. They give evidence not only for the ability to define significant deviations from normal but also to explore potential underlying mechanisms and therapeutic interventions.

A large body of evidence points to reduced BDNF delivery from cortical neurons as contributing to striatal degeneration in HD (Baydyuk and Xu, 2014; Gauthier et al., 2004; Su et al., 2014; Yu et al., 2018; Zuccato and Cattaneo, 2007, 2009). Using BACHD as a model system, and microfluidic chambers in which we created a cortico-striatal network, we previously showed that axonal transport of BDNF, both anterograde and retrograde, was markedly reduced (Zhao et al., 2016). The impact of reduced anterograde BDNF transport for the trophic status of BACHD striatal neurons was assessed by adding BDNF to the striatal neuron compartment, an intervention that prevented atrophy. These data raised the possibility of a presynaptic defect in BDNF delivery. Given a role for BDNF in regulating synapse formation and function (refs), we hypothesized that BACHD cortical neurons may demonstrate synaptic defects. Herein we addressed the synapse formation and function in BACHD cortical neuron cultures as compared to controls and asked if any defects discovered could be linked to changes in BDNF release.

We established long-term cultures of cortical neurons, extending observations as long as DIV35, and measured synapse formation and maintenance in WT as well as BACHD and ΔN17-BACHD cortical neurons. Our results demonstrated that cortical neurons from both mouse HD models were able to form functional synapses at DIV14. Thereafter, however, as compared to WT neurons BACHD neurons showed progressive reductions in synapse number and activity. These findings were consistent with our expectations. Inconsistent with our expectations were the findings in ΔN17-BACHD cultures. We expected to see regressive changes in synapses more dramatic than in BACHD mice because studies in the ΔN17-BACHD model documented more pronounced disease-linked phenotypes, including deficits in motor function(Gu et al., 2015). However, the opposite was found. At DIV 21 and later synapse number in ΔN17-BACHD cultures exceeded those in BACHD cultures. Other synaptic parameters in ΔN17-BACHD cultures also distinguished them from BACHD. Overall, synaptic measures in ΔN17-BACHD cultures were much more like WT cultures. Indeed, by DIV28 the number of synapses in ΔN17-BACHD cultures exceeded those in WT cultures. Taken together, we found dysregulated synaptic homeostasis in both the BACHD and ΔN17-BACHD cortical cultures. The findings for differences in BACHD and ΔN17-BACHD suggest that synapse structure is sensitive to the actions of mutant Htt proteins.

The increase in synaptic formation in ΔN17-BACHD neurons relative to both BACHD and WT neurons is intriguing. Several possibilities could explain the differences: 1) due to the absence of N17, ΔN17-mHTT accumulate in the nucleus (Gu et al., 2015)and therein could induce increased expression of genes regulating synapse maintenance. Attenuation of synaptically relevant transcriptional programs has been reported to follow maturation at DIV12-14 (Kavalali et al., 1999). It is possible that the presence of ΔN17-mHTT in the nucleus interferes with this regulatory process leading to excessive synthesis of synaptic proteins leading to anomalous new synapse formation in ΔN17-BACHD neurons. 2) Alternatively, the balance between the formation of new synapses and disappearance of existing synapses may be disrupted in favor of the formation of new synapses in the ΔN17-BACHD neurons. 3) It is also possible that synaptic dysfunction ΔN17-BACHD neurons, a topic not examined herein, engages a compensatory response under which synapses are formed in excess. 4) Finally, the ΔN17-BACHD protein may induce local changes in protein trafficking that promote synapse formation. Future studies are needed to explore these interesting possibilities with the goal of further elucidating not only the actions of ΔN17-BACHD but also mHTT as regards synapses.

The only significant difference between BACHD and ΔN17-BACHD is that the first N-terminal 17 residues were deleted in the ΔN17-BACHD model. While both variants of mHTT exhibit neuronal toxicity and lead to disease manifestations, our analysis revealing the dichotomy regarding synapses points to the amino terminus of the mutant protein as significantly impacting synaptic structure and maintenance. The N-terminal 17 amino acids of the protein (N17) was shown to enhance mHtt aggregation (Tam et al., 2009; Thakur et al., 2009) and to promote amyloid fibril formation in vitro (Shen et al., 2016). N17 is required for interaction with chaperones and is marked by many post-translational modification sites that apparently mediate mHtt toxicity (Gu et al., 2009; Steffan et al., 2004; Steffan and Thompson, 2003; Thompson et al., 2009). In addition, N17 dictates the subcellular localization of mHTT Exon1 protein and controls its targeting to mitochondria and association with the endoplasmic reticulum (ER) and Golgi (Rockabrand et al., 2007). N17 also functions as a nuclear export sequence (Maiuri et al., 2013; Rockabrand et al., 2007; Steffan et al., 2004; Steffan and Thompson, 2003; Zheng et al., 2018). It is unclear if existing findings for the localization and function of the full-length mutant Htt protein are sufficient to explain synaptic deficits in the BACHD mouse, but it is tempting to speculate that local synaptic changes involving protein sorting and mitochondrial function play roles.

Our previous study demonstrating reduced anterograde axonal transport of BDNF suggested that BDNF release at synapses was impaired. To test this directly, we harvested conditioned media at DIV14 and DIV21 from both WT and BACHD cultures. We found that BDNF secretion from BACHD neurons was significantly reduced (Gu et al., 2009; Steffan and Thompson, 2003) at DIV21, but not at DIV14. These findings point to age-related changes whose timing coincides with the reduction of synapses in BACHD cultures at DIV21, but not earlier. Given this finding, we asked if addition of exogenous BDNF to BACHD neurons would rescue synaptic deficits. We found that treatment of BACHD neurons for either 7 or 14 days effectively prevented synaptic deficits, restoring values to the levels in WT neurons. We found that ApiCCT1 also modestly improved synapse number. While the mechanism for this effect was not explored herein, in earlier work we found evidence for ApiCCT1-mediated downregulation of mHTT in BACHD neurons together with increased BDNF release from cortical axons (Zhao et al., 2016). However, we note the plurality of mHTT actions. As one example, a comprehensive proteomic analysis showed mHtt-induced alterations of many cellular pathways involved in synaptic transmission and plasticity (Kaltenbach et al., 2007; Shirasaki et al., 2012). Given this, and the inherent complexities in synapse formation and function, a thorough investigations of how mHTT impacts the synapse and how ApiCCT1 acts to intercept pathogenetic mechanisms is needed.

Deficits in synaptic maintenance in BACHD neurons predicted functional consequences and raised the possibility that BDNF would also normalize these changes. We used MEA to measure neuronal activity. Consistent with synapse formation, BACHD neurons did not exhibit overt deficits in only one of twelve metrics at DIV14. However, significant changes in four of twelve metrics were detected in BACHD neurons at DIV28. BACHD cultures showed dysregulation of synaptic activity and reduced network activity and synchrony. The findings are consistent with those for synapse number and other synaptic metrics at DIV28. Significantly, functional deficits were effectively rescued by BDNF treatment, with normalization of all the deficits present in BACHD neurons and with enhancement of synaptic functions not impacted in BACHD. As for synapse numbers, ApiCCT1 was much less effective than BDNF. We conclude that BDNF robustly acted to enhance synaptic function in BACHD neurons. Indeed, it exerted effects on some measures in BACHD neurons that appear to have exceeded those in WT neurons. A caveat for the interpretation of BDNF effects on synapse structure and function is that the levels of BDNF used almost certainly exceed those present in vivo. Whether not restoring BDNF release to physiological levels in vivo would restore synapses is not known and deserves additional study.

How BDNF acted to restore synapses is an important question. Previous studies have shown that decreased BDNF-TrkB signaling led to reduction in the density of striatal dendritic spines in both the BACHD and the Q175 knock-in mouse models of HD (Plotkin et al., 2014). BDNF overexpression in the forebrain effectively restored dendritic spines density and morphology in striatal neurons of HD mouse model (Xie et al., 2010). Furthermore, immunohistochemical staining against pre-synaptic (VGLUT1) and post-synaptic (PSD95) markers showed that the decrease of cortico-striatal synapses was significantly improved by increased expression of BDNF in HD models (Giralt et al., 2011). We envision that one effect of BDNF in restoring synapses in BACHD neurons is through enhancing the levels of synaptic proteins, as demonstrated by Giralt and colleagues.

In conclusion, in vitro studies exploring the BACHD and ΔN17-BACHD models of HD demonstrated evidence of significant synaptic deficits in the former and synaptic dysregulation in both. BACHD synaptic deficits were correlated with reduced release of BDNF and addition of BDNF to BACHD cultures restored and even enhanced synapse structure and function. The culture paradigm employed can be used to further explore HD synaptic pathogenesis and treatments to intercept synaptic pathology.

## Acknowledgements

The study is supported by NIH 5P01NS092525 (CW, WCM, JF, LMT) a gift from the Beckman Laser Institute Foundation, Air Force Office of Scientific Research under award number FA9550-17-1-0193 (CC, LS). We thank for technical assistances of Ruinan Shen, Savannah Fang, Kalina Wiatr, Simone Jetha. We also thank the Sue and Bill ross Stem Cell Center for use of MEA.

## References

The Huntington’s Disease Collaborative Research Group (1993). A novel gene containing a trinucleotide repeat that is expanded and unstable on Huntington’s disease chromosomes. Cell 72, 971–983.

Baquet, Z.C., Gorski, J.A., and Jones, K.R. (2004). Early striatal dendrite deficits followed by neuron loss with advanced age in the absence of anterograde cortical brain-derived neurotrophic factor. J Neurosci 24, 4250–4258.

Baydyuk, M., and Xu, B. (2014). BDNF signaling and survival of striatal neurons. Frontiers in Cellular Neuroscience 8.

Benchoua, A., Trioulier, Y., Zala, D., Gaillard, M.C., Lefort, N., Dufour, N., Saudou, F., Elalouf, J.M., Hirsch, E., Hantraye, P., et al. (2006). Involvement of mitochondrial complex II defects in neuronal death produced by N-terminus fragment of mutated huntingtin. Mol Biol Cell 17, 1652–1663.

Bennett, E.J., Shaler, T.A., Woodman, B., Ryu, K.Y., Zaitseva, T.S., Becker, C.H., Bates, G.P., Schulman, H., and Kopito, R.R. (2007). Global changes to the ubiquitin system in Huntington’s disease. Nature 448, 704–708.

Blumenstock, S., and Dudanova, I. (2020). Cortical and Striatal Circuits in Huntington’s Disease. Frontiers in Neuroscience 14.

Browne, S.E. (2008). Mitochondria and Huntington’s disease pathogenesis: insight from genetic and chemical models. Ann N Y Acad Sci 1147, 358–382.

Carmo, C., Naia, L., Lopes, C., and Rego, A.C. (2018). Mitochondrial Dysfunction in Huntington’s Disease. Adv Exp Med Biol 1049, 59–83.

Caron, N.S., Dorsey, E.R., and Hayden, M.R. (2018). Therapeutic approaches to Huntington disease: from the bench to the clinic. Nature Reviews Drug Discovery 17, 729–750.

Chen, F., Ardalan, M., Elfving, B., Wegener, G., Madsen, T.M., and Nyengaard, J.R. (2018a). Mitochondria Are Critical for BDNF-Mediated Synaptic and Vascular Plasticity of Hippocampus following Repeated Electroconvulsive Seizures. Int J Neuropsychopharmacol 21, 291–304.

Chen, X.-Q., Fang, F., Florio, J.B., Rockenstein, E., Masliah, E., Mobley, W.C., Rissman, R.A., and Wu, C. (2018b). T-complex protein 1-ring complex enhances retrograde axonal transport by modulating tau phosphorylation. Traffic 19, 840–853.

Conner, J.M., Lauterborn, J.C., Yan, Q., Gall, C.M., and Varon, S. (1997). Distribution of brain-derived neurotrophic factor (BDNF) protein and mRNA in the normal adult rat CNS: evidence for anterograde axonal transport. J Neurosci 17, 2295–2313.

Cummings, D.M., Milnerwood, A.J., Dallerac, G.M., Vatsavayai, S.C., Hirst, M.C., and Murphy, K.P. (2007). Abnormal cortical synaptic plasticity in a mouse model of Huntington’s disease. Brain Res Bull 72, 103–107.

Dallerac, G.M., Vatsavayai, S.C., Cummings, D.M., Milnerwood, A.J., Peddie, C.J., Evans, K.A., Walters, S.W., Rezaie, P., Hirst, M.C., and Murphy, K.P. (2011). Impaired long-term potentiation in the prefrontal cortex of Huntington’s disease mouse models: rescue by D1 dopamine receptor activation. Neurodegener Dis 8, 230–239.

Damiano, M., Diguet, E., Malgorn, C., D’Aurelio, M., Galvan, L., Petit, F., Benhaim, L., Guillermier, M., Houitte, D., Dufour, N., et al. (2013). A role of mitochondrial complex II defects in genetic models of Huntington’s disease expressing N-terminal fragments of mutant huntingtin. Hum Mol Genet 22, 3869–3882.

Daniele, J.R., Heydari, K., Arriaga, E.A., and Dillin, A. (2016). Identification and Characterization of Mitochondrial Subtypes in Caenorhabditis elegans via Analysis of Individual Mitochondria by Flow Cytometry. Anal Chem 88, 6309–6316.

Duffey, C.A., Nicholls, S., Mantilla, C.B., Fogarty, M.J., and Sieck, G.C. (2018). Effect of BDNF on Mitochondrial Morphology and Protein Expression in NSC-34 Cells. The FASEB Journal 32, 743.746-743.746.

Fang, F., Yang, W., Florio, J.B., Rockenstein, E., Spencer, B., Orain, X.M., Dong, S.X., Li, H., Chen, X., Sung, K., et al. (2017). Synuclein impairs trafficking and signaling of BDNF in a mouse model of Parkinson’s disease. Scientific Reports 7, 3868.

Finkbeiner, S. (2011). Huntington’s Disease. Cold Spring Harb Perspect Biol 3.

Frank, S. (2014). Treatment of Huntington’s disease. Neurotherapeutics 11, 153-160.

Frydman, J. (2001). Folding of newly translated proteins in vivo: the role of molecular chaperones. Annu Rev Biochem 70, 603–647.

Frydman, J., Nimmesgern, E., Erdjument-Bromage, H., Wall, J.S., Tempst, P., and Hartl, F.U. (1992). Function in protein folding of TRiC, a cytosolic ring complex containing TCP-1 and structurally related subunits. EMBO J 11, 4767–4778.

Gauthier, L.R., Charrin, B.C., Borrell-Pages, M., Dompierre, J.P., Rangone, H., Cordelieres, F.P., De Mey, J., MacDonald, M.E., Lessmann, V., Humbert, S., et al. (2004). Huntingtin controls neurotrophic support and survival of neurons by enhancing BDNF vesicular transport along microtubules. Cell 118, 127–138.

Gestaut, D., Roh, S.H., Ma, B., Pintilie, G., Joachimiak, L.A., Leitner, A., Walzthoeni, T., Aebersold, R., Chiu, W., and Frydman, J. (2019). The Chaperonin TRiC/CCT Associates with Prefoldin through a Conserved Electrostatic Interface Essential for Cellular Proteostasis. Cell 177, 751–765.e715.

Giralt, A., Carreton, O., Lao-Peregrin, C., Martin, E.D., and Alberch, J. (2011). Conditional BDNF release under pathological conditions improves Huntington’s disease pathology by delaying neuronal dysfunction. Mol Neurodegener 6, 71.

Gorski, J.A., Zeiler, S.R., Tamowski, S., and Jones, K.R. (2003). Brain-derived neurotrophic factor is required for the maintenance of cortical dendrites. J Neurosci 23, 6856–6865.

Gray, M., Shirasaki, D.I., Cepeda, C., Andre, V.M., Wilburn, B., Lu, X.H., Tao, J., Yamazaki, I., Li, S.H., Sun, Y.E., et al. (2008). Full-length human mutant huntingtin with a stable polyglutamine repeat can elicit progressive and selective neuropathogenesis in BACHD mice. J Neurosci 28, 6182–6195.

Gu, X., Cantle, J.P., Greiner, E.R., Lee, C.Y., Barth, A.M., Gao, F., Park, C.S., Zhang, Z., Sandoval-Miller, S., Zhang, R.L., et al. (2015). N17 Modifies mutant Huntingtin nuclear pathogenesis and severity of disease in HD BAC transgenic mice. Neuron 85, 726–741.

Gu, X., Greiner, E.R., Mishra, R., Kodali, R., Osmand, A., Finkbeiner, S., Steffan, J.S., Thompson, L.M., Wetzel, R., and Yang, X.W. (2009). Serines 13 and 16 are critical determinants of full-length human mutant huntingtin induced disease pathogenesis in HD mice. Neuron 64, 828–840.

Guedes-Dias, P., Pinho, B.R., Soares, T.R., de Proenca, J., Duchen, M.R., and Oliveira, J.M. (2016). Mitochondrial dynamics and quality control in Huntington’s disease. Neurobiol Dis 90, 51–57.

Guo, X., Sun, X., Hu, D., Wang, Y.J., Fujioka, H., Vyas, R., Chakrapani, S., Joshi, A.U., Luo, Y., Mochly-Rosen, D., et al. (2016). VCP recruitment to mitochondria causes mitophagy impairment and neurodegeneration in models of Huntington’s disease. Nat Commun 7, 12646.

Hofer, M., Pagliusi, S.R., Hohn, A., Leibrock, J., and Barde, Y.A. (1990). Regional distribution of brain-derived neurotrophic factor mRNA in the adult mouse brain. EMBO J 9, 2459–2464.

Hong, Y., Zhao, T., Li, X.-J., and Li, S. (2016). Mutant Huntingtin Impairs BDNF Release from Astrocytes by Disrupting Conversion of Rab3a-GTP into Rab3a-GDP. The Journal of Neuroscience 36, 8790.

Kabir, M.A., Uddin, W., Narayanan, A., Reddy, P.K., Jairajpuri, M.A., Sherman, F., and Ahmad, Z. (2011). Functional Subunits of Eukaryotic Chaperonin CCT/TRiC in Protein Folding. J Amino Acids 2011, 843206.

Kalisman, N., Adams, C.M., and Levitt, M. (2012). Subunit order of eukaryotic TRiC/CCT chaperonin by cross-linking, mass spectrometry, and combinatorial homology modeling. Proc Natl Acad Sci U S A 109, 2884–2889.

Kaltenbach, L.S., Romero, E., Becklin, R.R., Chettier, R., Bell, R., Phansalkar, A., Strand, A., Torcassi, C., Savage, J., Hurlburt, A., et al. (2007). Huntingtin interacting proteins are genetic modifiers of neurodegeneration. PLoS Genet 3, e82.

Kavalali, E.T., Klingauf, J., and Tsien, R.W. (1999). Activity-dependent regulation of synaptic clustering in a hippocampal culture system. Proceedings of the National Academy of Sciences 96, 12893.

Kitamura, A., Kubota, H., Pack, C.G., Matsumoto, G., Hirayama, S., Takahashi, Y., Kimura, H., Kinjo, M., Morimoto, R.I., and Nagata, K. (2006). Cytosolic chaperonin prevents polyglutamine toxicity with altering the aggregation state. Nat Cell Biol 8, 1163–1170.

Klapstein, G.J., Fisher, R.S., Zanjani, H., Cepeda, C., Jokel, E.S., Chesselet, M.F., and Levine, M.S. (2001). Electrophysiological and morphological changes in striatal spiny neurons in R6/2 Huntington’s disease transgenic mice. J Neurophysiol 86, 2667–2677.

Kovalenko, M., Milnerwood, A., Giordano, J., St Claire, J., Guide, J.R., Stromberg, M., Gillis, T., Sapp, E., DiFiglia, M., MacDonald, M.E., et al. (2018). HttQ111/+ Huntington’s Disease Knock-in Mice Exhibit Brain Region-Specific Morphological Changes and Synaptic Dysfunction. J Huntingtons Dis 7, 17–33.

Lane, R.M., Smith, A., Baumann, T., Gleichmann, M., Norris, D., Bennett, C.F., and Kordasiewicz, H. (2018). Translating Antisense Technology into a Treatment for Huntington’s Disease. Methods Mol Biol 1780, 497–523.

Leitman, J., Ulrich Hartl, F., and Lederkremer, G.Z. (2013). Soluble forms of polyQ-expanded huntingtin rather than large aggregates cause endoplasmic reticulum stress. Nat Commun 4, 2753.

Leitner, A., Joachimiak, L.A., Bracher, A., Monkemeyer, L., Walzthoeni, T., Chen, B., Pechmann, S., Holmes, S., Cong, Y., Ma, B., et al. (2012). The molecular architecture of the eukaryotic chaperonin TRiC/CCT. Structure 20, 814–825.

Li, X., Valencia, A., Sapp, E., Masso, N., Alexander, J., Reeves, P., Kegel, K.B., Aronin, N., and Difiglia, M. (2010). Aberrant Rab11-dependent trafficking of the neuronal glutamate transporter EAAC1 causes oxidative stress and cell death in Huntington’s disease. J Neurosci 30, 4552–4561.

Lindsay, R.M., Wiegand, S.J., Altar, C.A., and DiStefano, P.S. (1994). Neurotrophic factors: from molecule to man. Trends Neurosci 17, 182–190.

Lu, B., Nagappan, G., and Lu, Y. (2014). BDNF and synaptic plasticity, cognitive function, and dysfunction. Handb Exp Pharmacol 220, 223–250.

Maisonpierre, P.C., Belluscio, L., Friedman, B., Alderson, R.F., Wiegand, S.J., Furth, M.E., Lindsay, R.M., and Yancopoulos, G.D. (1990). NT-3, BDNF, and NGF in the developing rat nervous system: parallel as well as reciprocal patterns of expression. Neuron 5, 501–509.

Maiuri, T., Woloshansky, T., Xia, J., and Truant, R. (2013). The huntingtin N17 domain is a multifunctional CRM1 and Ran-dependent nuclear and cilial export signal. Hum Mol Genet 22, 1383–1394.

Mangiarini, L., Sathasivam, K., Seller, M., Cozens, B., Harper, A., Hetherington, C., Lawton, M., Trottier, Y., Lehrach, H., Davies, S.W., et al. (1996). Exon 1 of the HD gene with an expanded CAG repeat is sufficient to cause a progressive neurological phenotype in transgenic mice. Cell 87, 493–506.

Markham, A., Cameron, I., Franklin, P., and Spedding, M. (2004). BDNF increases rat brain mitochondrial respiratory coupling at complex I, but not complex II. Eur J Neurosci 20, 1189–1196.

McColgan, P., and Tabrizi, S.J. (2018). Huntington’s disease: a clinical review. Eur J Neurol 25, 24–34.

Milnerwood, A.J., Cummings, D.M., Dallerac, G.M., Brown, J.Y., Vatsavayai, S.C., Hirst, M.C., Rezaie, P., and Murphy, K.P. (2006). Early development of aberrant synaptic plasticity in a mouse model of Huntington’s disease. Hum Mol Genet 15, 1690–1703.

Milnerwood, A.J., and Raymond, L.A. (2007). Corticostriatal synaptic function in mouse models of Huntington’s disease: early effects of huntingtin repeat length and protein load. J Physiol 585, 817–831.

Murphy, K.P., Carter, R.J., Lione, L.A., Mangiarini, L., Mahal, A., Bates, G.P., Dunnett, S.B., and Morton, A.J. (2000). Abnormal synaptic plasticity and impaired spatial cognition in mice transgenic for exon 1 of the human Huntington’s disease mutation. J Neurosci 20, 5115–5123.

Nithianantharajah, J., and Hannan, A.J. (2013). Dysregulation of synaptic proteins, dendritic spine abnormalities and pathological plasticity of synapses as experience-dependent mediators of cognitive and psychiatric symptoms in Huntington’s disease. Neuroscience 251, 66–74.

Nollen, E.A., Garcia, S.M., van Haaften, G., Kim, S., Chavez, A., Morimoto, R.I., and Plasterk, R.H. (2004). Genome-wide RNA interference screen identifies previously undescribed regulators of polyglutamine aggregation. Proc Natl Acad Sci U S A 101, 6403–6408.

Nosyreva, E., Szabla, K., Autry, A.E., Ryazanov, A.G., Monteggia, L.M., and Kavalali, E.T. (2013). Acute Suppression of Spontaneous Neurotransmission Drives Synaptic Potentiation. The Journal of Neuroscience 33, 6990.

Park, H., and Poo, M.M. (2013). Neurotrophin regulation of neural circuit development and function. Nat Rev Neurosci 14, 7–23.

Plotkin, J.L., Day, M., Peterson, J.D., Xie, Z., Kress, G.J., Rafalovich, I., Kondapalli, J., Gertler, T.S., Flajolet, M., Greengard, P., et al. (2014). Impaired TrkB receptor signaling underlies corticostriatal dysfunction in Huntington’s disease. Neuron 83, 178–188.

Puigdellivol, M., Cherubini, M., Brito, V., Giralt, A., Suelves, N., Ballesteros, J., Zamora-Moratalla, A., Martin, E.D., Eipper, B.A., Alberch, J., et al. (2015). A role for Kalirin-7 in corticostriatal synaptic dysfunction in Huntington’s disease. Hum Mol Genet 24, 7265–7285.

Reddy, P.H., Mao, P., and Manczak, M. (2009). Mitochondrial structural and functional dynamics in Huntington’s disease. Brain Res Rev 61, 33–48.

Rockabrand, E., Slepko, N., Pantalone, A., Nukala, V.N., Kazantsev, A., Marsh, J.L., Sullivan, P.G., Steffan, J.S., Sensi, S.L., and Thompson, L.M. (2007). The first 17 amino acids of Huntingtin modulate its sub-cellular localization, aggregation and effects on calcium homeostasis. Hum Mol Genet 16, 61–77.

Rodinova, M., Krizova, J., Stufkova, H., Bohuslavova, B., Askeland, G., Dosoudilova, Z., Juhas, S., Juhasova, J., Ellederova, Z., Zeman, J., et al. (2019). Deterioration of mitochondrial bioenergetics and ultrastructure impairment in skeletal muscle of a transgenic minipig model in the early stages of Huntington’s disease. Dis Model Mech 12.

Roh, S.H., Kasembeli, M., Bakthavatsalam, D., Chiu, W., and Tweardy, D.J. (2015). Contribution of the Type II Chaperonin, TRiC/CCT, to Oncogenesis. Int J Mol Sci 16, 26706–26720.

Santos, A.R., Comprido, D., and Duarte, C.B. (2010). Regulation of local translation at the synapse by BDNF. Prog Neurobiol 92, 505–516.

Saudou, F., and Humbert, S. (2016). The Biology of Huntingtin. Neuron 89, 910–926.

Sharp, A.H., Loev, S.J., Schilling, G., Li, S.H., Li, X.J., Bao, J., Wagster, M.V., Kotzuk, J.A., Steiner, J.P., Lo, A., et al. (1995). Widespread expression of Huntington’s disease gene (IT15) protein product. Neuron 14, 1065–1074.

Shen, K., Calamini, B., Fauerbach, J.A., Ma, B., Shahmoradian, S.H., Serrano Lachapel, I.L., Chiu, W., Lo, D.C., and Frydman, J. (2016). Control of the structural landscape and neuronal proteotoxicity of mutant Huntingtin by domains flanking the polyQ tract. eLife 5, e18065.

Shirasaki, D.I., Greiner, E.R., Al-Ramahi, I., Gray, M., Boontheung, P., Geschwind, D.H., Botas, J., Coppola, G., Horvath, S., Loo, J.A., et al. (2012). Network organization of the huntingtin proteomic interactome in mammalian brain. Neuron 75, 41–57.

Shirendeb, U.P., Calkins, M.J., Manczak, M., Anekonda, V., Dufour, B., McBride, J.L., Mao, P., and Reddy, P.H. (2012). Mutant huntingtin’s interaction with mitochondrial protein Drp1 impairs mitochondrial biogenesis and causes defective axonal transport and synaptic degeneration in Huntington’s disease. Hum Mol Genet 21, 406–420.

Smith-Dijak, A.I., Sepers, M.D., and Raymond, L.A. (2019). Alterations in synaptic function and plasticity in Huntington disease. J Neurochem 150, 346–365.

Song, W., Chen, J., Petrilli, A., Liot, G., Klinglmayr, E., Zhou, Y., Poquiz, P., Tjong, J., Pouladi, M.A., Hayden, M.R., et al. (2011). Mutant huntingtin binds the mitochondrial fission GTPase dynamin-related protein-1 and increases its enzymatic activity. Nat Med 17, 377–382.

Sontag, E.M., Joachimiak, L.A., Tan, Z., Tomlinson, A., Housman, D.E., Glabe, C.G., Potkin, S.G., Frydman, J., and Thompson, L.M. (2013). Exogenous delivery of chaperonin subunit fragment ApiCCT1 modulates mutant Huntingtin cellular phenotypes. Proc Natl Acad Sci U S A 110, 3077–3082.

Steffan, J.S., Agrawal, N., Pallos, J., Rockabrand, E., Trotman, L.C., Slepko, N., Illes, K., Lukacsovich, T., Zhu, Y.Z., Cattaneo, E., et al. (2004). SUMO modification of Huntingtin and Huntington’s disease pathology. Science 304, 100–104.

Steffan, J.S., and Thompson, L.M. (2003). Targeting aggregation in the development of therapeutics for the treatment of Huntington’s disease and other polyglutamine repeat diseases. Expert Opin Ther Targets 7, 201–213.

Su, B., Ji, Y.-S., Sun, X.-l., Liu, X.-H., and Chen, Z.-Y. (2014). Brain-derived neurotrophic factor (BDNF)-induced mitochondrial motility arrest and presynaptic docking contribute to BDNF-enhanced synaptic transmission. J Biol Chem 289, 1213–1226.

Tam, S., Geller, R., Spiess, C., and Frydman, J. (2006). The chaperonin TRiC controls polyglutamine aggregation and toxicity through subunit-specific interactions. Nat Cell Biol 8, 1155–1162.

Tam, S., Spiess, C., Auyeung, W., Joachimiak, L., Chen, B., Poirier, M.A., and Frydman, J. (2009). The chaperonin TRiC blocks a huntingtin sequence element that promotes the conformational switch to aggregation. Nature Structural & Molecular Biology 16, 1279–1285.

Thakur, A.K., Jayaraman, M., Mishra, R., Thakur, M., Chellgren, V.M., L Byeon, I.-J., Anjum, D.H., Kodali, R., Creamer, T.P., Conway, J.F., et al. (2009). Polyglutamine disruption of the huntingtin exon 1 N terminus triggers a complex aggregation mechanism. Nature Structural & Molecular Biology 16, 380–389.

Thulasiraman, V., Yang, C.F., and Frydman, J. (1999). In vivo newly translated polypeptides are sequestered in a protected folding environment. EMBO J 18, 85–95.

Veldman, M.B., and Yang, X.W. (2018). Molecular insights into cortico-striatal miscommunications in Huntington’s disease. Curr Opin Neurobiol 48, 79–89.

Vonsattel, J.P., and DiFiglia, M. (1998). Huntington disease. J Neuropathol Exp Neurol 57, 369–384.

Walker, F.O. (2007). Huntington’s disease. Lancet 369, 218–228.

Xie, Y., Hayden, M.R., and Xu, B. (2010). BDNF overexpression in the forebrain rescues Huntington’s disease phenotypes in YAC128 mice. J Neurosci 30, 14708–14718.

Yablonska, S., Ganesan, V., Ferrando, L.M., Kim, J., Pyzel, A., Baranova, O.V., Khattar, N.K., Larkin, T.M., Baranov, S.V., Chen, N., et al. (2019). Mutant huntingtin disrupts mitochondrial proteostasis by interacting with TIM23. Proceedings of the National Academy of Sciences of the United States of America 116, 16593–16602.

Yan, Q., Rosenfeld, R.D., Matheson, C.R., Hawkins, N., Lopez, O.T., Bennett, L., and Welcher, A.A. (1997). Expression of brain-derived neurotrophic factor protein in the adult rat central nervous system. Neuroscience 78, 431–448.

Yoshii, A., and Constantine-Paton, M. (2010). Postsynaptic BDNF-TrkB signaling in synapse maturation, plasticity, and disease. Dev Neurobiol 70, 304–322.

Yu, C., Li, C.H., Chen, S., Yoo, H., Qin, X., and Park, H. (2018). Decreased BDNF Release in Cortical Neurons of a Knock-in Mouse Model of Huntington’s Disease. Scientific Reports 8, 16976.

Zhao, X., Chen, X.Q., Han, E., Hu, Y., Paik, P., Ding, Z., Overman, J., Lau, A.L., Shahmoradian, S.H., Chiu, W., et al. (2016). TRiC subunits enhance BDNF axonal transport and rescue striatal atrophy in Huntington’s disease. Proc Natl Acad Sci U S A 113, E5655–5664.

Zheng, J., Winderickx, J., Franssens, V., and Liu, B. (2018). A Mitochondria-Associated Oxidative Stress Perspective on Huntington’s Disease. Front Mol Neurosci 11, 329.

Zheng, Z., Li, A., Holmes, B.B., Marasa, J.C., and Diamond, M.I. (2013). An N-terminal nuclear export signal regulates trafficking and aggregation of Huntingtin (Htt) protein exon 1. J Biol Chem 288, 6063–6071.

Zuccato, C., and Cattaneo, E. (2007). Role of brain-derived neurotrophic factor in Huntington’s disease. Prog Neurobiol 81, 294–330.

## References

Barry, J., Bui, M.T., Levine, M.S., and Cepeda, C. (2022). Synaptic pathology in Huntington’s disease: Beyond the corticostriatal pathway. Neurobiology of Disease 162, 105574.

Cepeda, C., and Levine, M.S. (2022). Synaptic Dysfunction in Huntington’s Disease: Lessons from Genetic Animal Models. The Neuroscientist 28, 20–40.

Chen, Y., Fu, A.K., and Ip, N.Y. (2019). Synaptic dysfunction in Alzheimer’s disease: Mechanisms and therapeutic strategies. Pharmacology & therapeutics 195, 186–198.

Cotterill, E., Charlesworth, P., Thomas, C.W., Paulsen, O., and Eglen, S.J. (2016). A comparison of computational methods for detecting bursts in neuronal spike trains and their application to human stem cell-derived neuronal networks. J Neurophysiol 116, 306–321.

Ferrer, I., Goutan, E., Marín, C., Rey, M.J., and Ribalta, T. (2000). Brain-derived neurotrophic factor in Huntington disease. Brain Res 866, 257–261.

Finkbeiner, S. (2011). Huntington’s Disease. Cold Spring Harb Perspect Biol 3.

Frank, S. (2014). Treatment of Huntington’s disease. Neurotherapeutics 11, 153-160.

Group, T.H.s.D.C.R. (1993). A novel gene containing a trinucleotide repeat that is expanded and unstable on Huntington’s disease chromosomes.. Cell 72, 971–983.

Kolodziejczyk, K., Parsons, M.P., Southwell, A.L., Hayden, M.R., and Raymond, L.A. (2014). Striatal synaptic dysfunction and hippocampal plasticity deficits in the Hu97/18 mouse model of Huntington disease. PLoS One 9, e94562.

Komatsu, H. (2021). Innovative Therapeutic Approaches for Huntington’s Disease: From Nucleic Acids to GPCR-Targeting Small Molecules. Frontiers in Cellular Neuroscience 15.

Lepeta, K., Lourenco, M.V., Schweitzer, B.C., Martino Adami, P.V., Banerjee, P., Catuara-Solarz, S., de La Fuente Revenga, M., Guillem, A.M., Haidar, M., Ijomone, O.M., et al. (2016). Synaptopathies: synaptic dysfunction in neurological disorders - A review from students to students. Journal of neurochemistry 138, 785–805.

Li, J.-Y., Plomann, M., and Brundin, P. (2003). Huntington’s disease: a synaptopathy? Trends in Molecular Medicine 9, 414–420.

Maity, S., Komal, P., Kumar, V., Saxena, A., Tungekar, A., and Chandrasekar, V. (2022). Impact of ER stress and ER-mitochondrial crosstalk in huntington’s disease. International Journal of Molecular Sciences 23, 780.

Morton, A., Faull, R., and Edwardson, J. (2001). Abnormalities in the synaptic vesicle fusion machinery in Huntington’s disease. Brain research bulletin 56, 111–117.

Novak, M.J.U., and Tabrizi, S.J. (2011). Huntington’s Disease: Clinical Presentation and Treatment. In International Review of Neurobiology, J. Brotchie, E. Bezard, and P. Jenner, eds. (Academic Press), pp. 297–323.

Pan, L., and Feigin, A. (2021). Huntington’s disease: New frontiers in therapeutics. Current Neurology and Neuroscience Reports 21, 1–9.

Paul, G., Cardinale, J., and Sbalzarini, I.F. (2013). Coupling Image Restoration and Segmentation: A Generalized Linear Model/Bregman Perspective. International Journal of Computer Vision 104, 69–93.

Rizk, A., Paul, G., Incardona, P., Bugarski, M., Mansouri, M., Niemann, A., Ziegler, U., Berger, P., and Sbalzarini, I.F. (2014). Segmentation and quantification of subcellular structures in fluorescence microscopy images using Squassh. Nat Protoc 9, 586–596.

Roos, R.A.C. (2010). Huntington’s disease: a clinical review. Orphanet J Rare Dis 5, 40–40.

Sahl, S.J., Lau, L., Vonk, W.I., Weiss, L.E., Frydman, J., and Moerner, W.E. (2016). Delayed emergence of subdiffraction-sized mutant huntingtin fibrils following inclusion body formation. Q Rev Biophys 49, e2.

Schirinzi, T., Madeo, G., Martella, G., Maltese, M., Picconi, B., Calabresi, P., and Pisani, A. (2016). Early synaptic dysfunction in Parkinson’s disease: insights from animal models. Movement Disorders 31, 802–813.

Selkoe, D.J. (2002). Alzheimer’s disease is a synaptic failure. Science 298, 789–791.

Su, B., Ji, Y.-S., Sun, X.-l., Liu, X.-H., and Chen, Z.-Y. (2014). Brain-derived neurotrophic factor (BDNF)-induced mitochondrial motility arrest and presynaptic docking contribute to BDNF-enhanced synaptic transmission. The Journal of biological chemistry 289, 1213–1226.

Subramanian, J., and Tremblay, M.-È. (2021). Synaptic Loss and Neurodegeneration (Frontiers Media SA), pp. 681029.

Taoufik, E., Kouroupi, G., Zygogianni, O., and Matsas, R. (2018). Synaptic dysfunction in neurodegenerative and neurodevelopmental diseases: an overview of induced pluripotent stem-cell-based disease models. Open Biol 8, 180138.

Thompson, L.M., Aiken, C.T., Kaltenbach, L.S., Agrawal, N., Illes, K., Khoshnan, A., Martinez-Vincente, M., Arrasate, M., O’Rourke, J.G., Khashwji, H., et al. (2009). IKK phosphorylates Huntingtin and targets it for degradation by the proteasome and lysosome. J Cell Biol 187, 1083–1099.

Vonsattel, J.P., and DiFiglia, M. (1998). Huntington disease. J Neuropathol Exp Neurol 57, 369–384.

Walker, F.O. (2007). Huntington’s disease. Lancet 369, 218–228.

Zuccato, C., and Cattaneo, E. (2009). Brain-derived neurotrophic factor in neurodegenerative diseases. Nat Rev Neurol 5, 311–322.

