## Supplemental Figures for "BDNF and TRiC-inspired Reagents Rescue Cortical Synaptic Deficits in a Mouse Model of Huntington’s Disease"

**Figure S1. Western analysis of changes in synaptic proteins.**
WT and BACHD neurons were cultured and protein lysates were collected in RIPA buffer at DIV21 and DIV28. Protein lysates were analyzed by SDS-PAGE/immunoblotting with specific antibodies as indicated. Beta actin was used as a loading control. Exposures within linear range were quantitated using BioRad Image Lab 6.0 and are normalized against beta actin. Significance tests are performed in Prism (unpaired t-test). * $p<0.05$; ** $p<0.01$; *** $p<0.001$.

**Figure S2. Measurement of secreted BDNF in conditioned media from cultured neurons**
E18 cortical neurons from WT and BACHD were dissected, cultured on PLL-coated 12-well plates and maintained as described in Materials and Methods. Conditioned media were collected at DIVs and the amounts of BDNF were measured by ELISA as described in the Materials and Methods. Corresponding cell lysates were collected with total proteins measured using BCA. Secreted BDNF was normalized against the respective protein content of each sample. (A) Standard curve of the first measurement. (B) Comparison of BDNF secretion in conditioned media in WT and BACHD cortical cultures at DIV14 and DIV21. (C) Standard curve of the second measurement. (D) Same comparison as (B). Results are shown as mean ± SEM. Significance analysis was carried out using Prism. Statistical significances were calculated by unpaired Student's t test. n.s.= non significance. All p values are shown in the graphs.

**Figure S3. Rescuing effect of BDNF on BACHD synaptic activity at DIV28.**
After completion of recording at DIV14, WT and BACHD neuronal cultures were treated with BDNF (50 ng/ml) or Vehicle. Media were changed every 48 hrs and a final recording was performed as above at DIV 28. MEA recording was performed as in Fig. 9, 10. A: WT neurons treated with vehicle; B: BACHD neurons treated with vehicle; C: WT neurons treated with BDNF; D. BACHD neurons treated with BDNF.

Fig S1. Western analsysis of synaptic proteins in cortical cultures from WT and BACHD

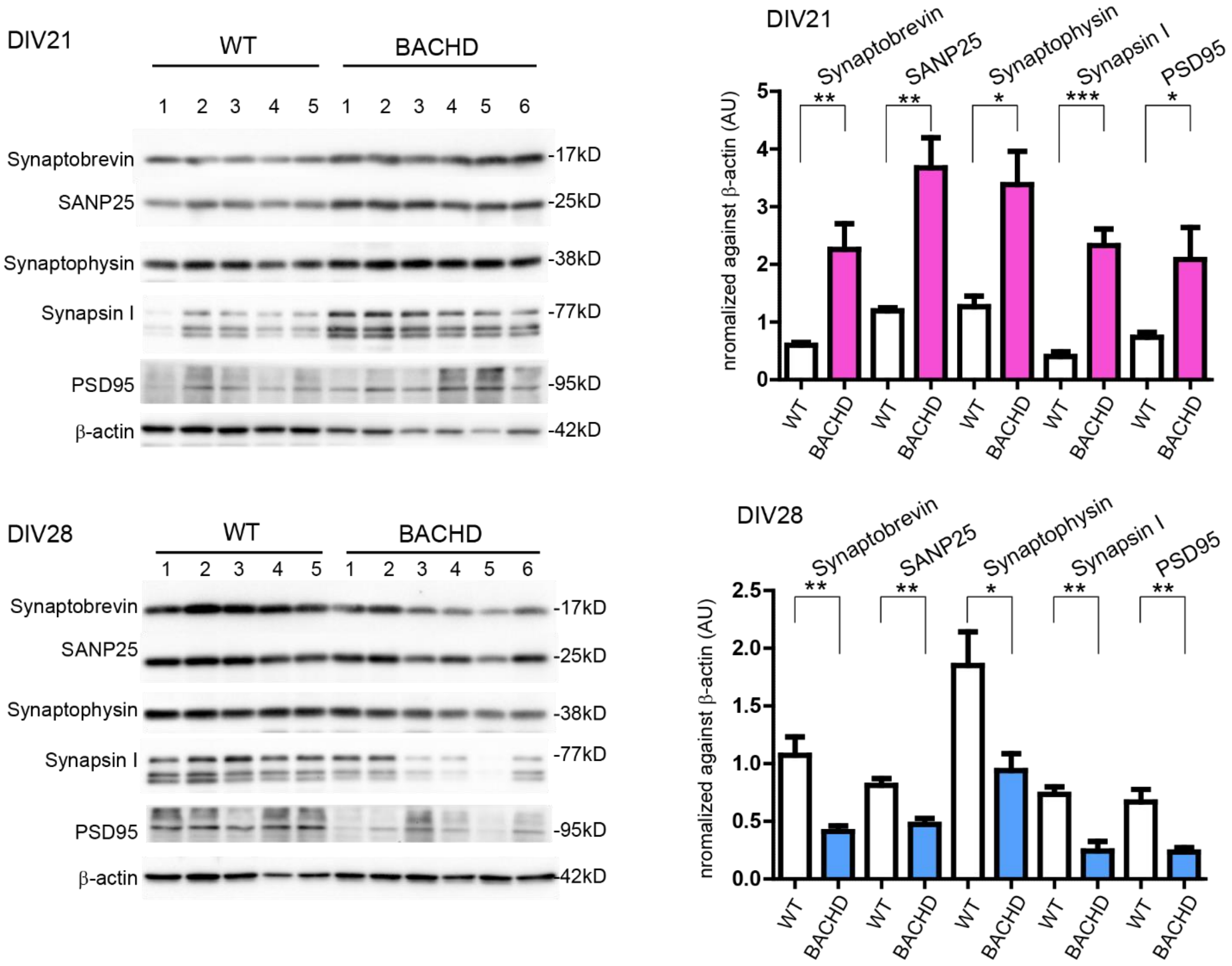

**Fig S2. Reduced BDNF secretion in BACHD cortical neurons (Gu et al., 2022)**

Cortical neuronal BDNF secretion (DIV14 vs 21)

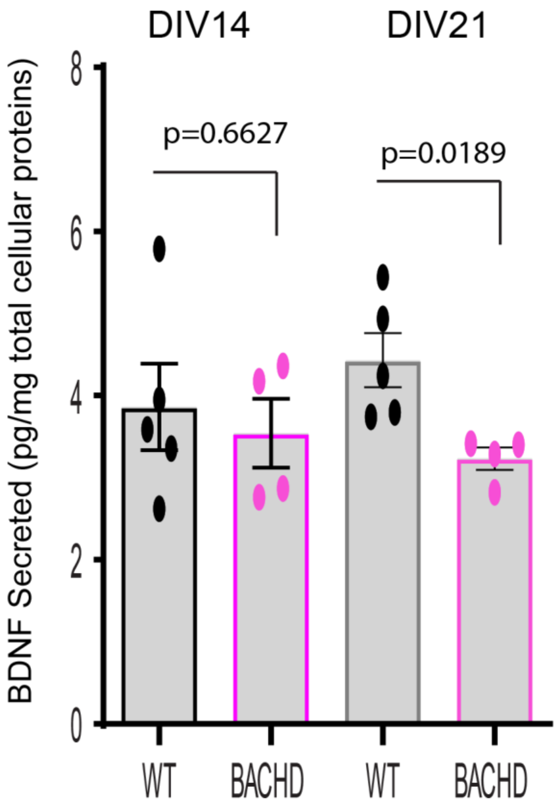

Fig S3. Effects of BDNF in WT and BACHD neurons

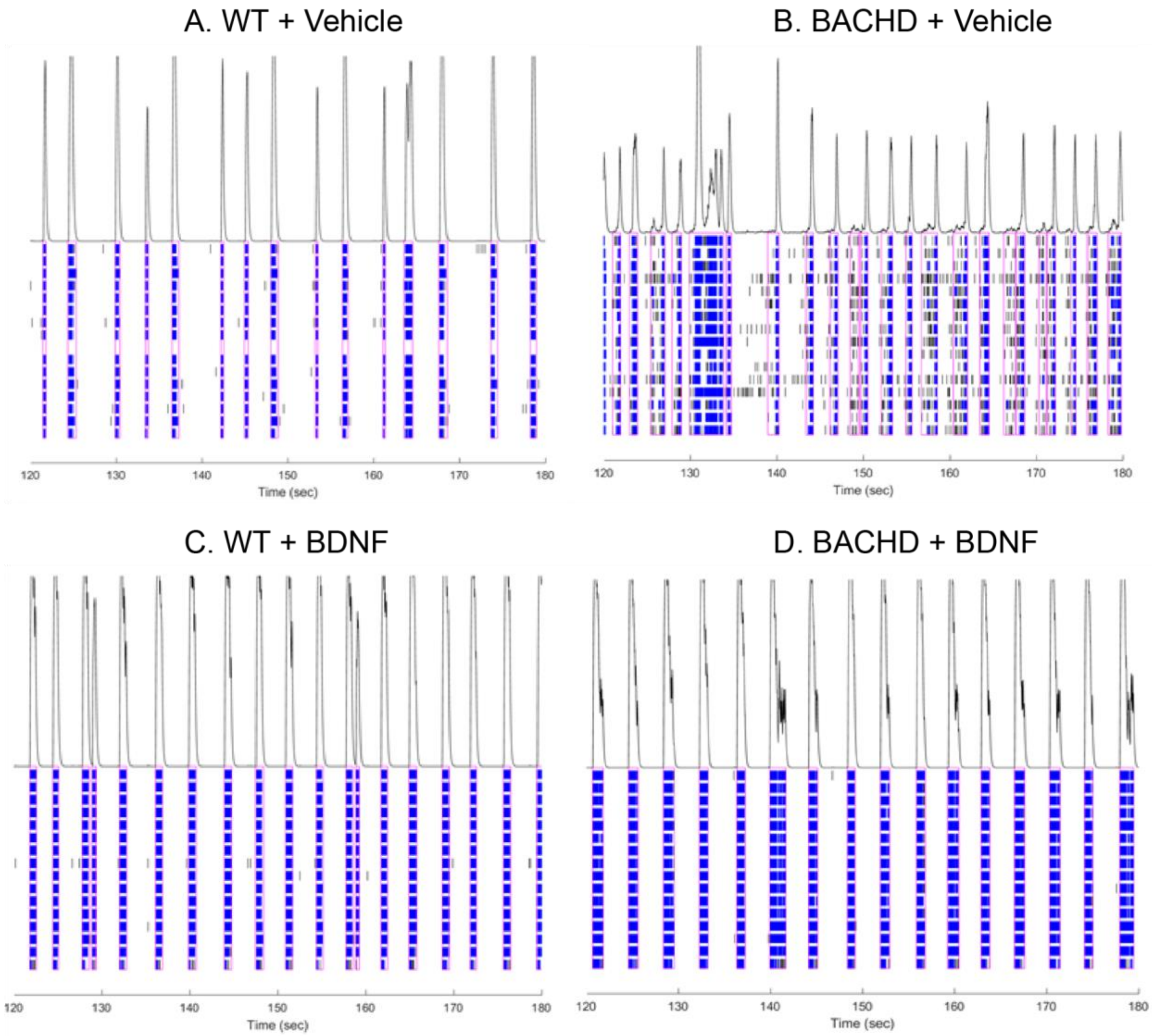
